# Structural and dynamic embedding of the mouse functional connectome revealed by functional ultrasound imaging (fUSI)

**DOI:** 10.64898/2026.02.05.704055

**Authors:** Chiara Pepe, Jean-Charles Mariani, Mila Urosevic, Silvia Gini, Alexia Stuefer, Fabio Ricci, Alberto Galbusera, Giuliano Iurilli, Alessandro Gozzi

## Abstract

Functional ultrasound imaging (fUSI) is an emerging hemodynamic neuroimaging modality whose potential for connectome-scale network mapping remains largely untested. Here, we establish a non-invasive transcranial fUSI protocol and an fMRI-inspired preprocessing framework that enable robust resting-state functional connectomics in the mouse. We show that fUSI resolves canonical brain-wide cortical and subcortical networks with high spatial concordance to fMRI, including a default-mode (DMN) and a laterocortical network. Notably, fUSI networks are robustly embedded within the structural connectome, with structure-function coupling being parsimoniously described by four dominant axes that differentially relate functional systems to known anatomical substrates. We also show that, beyond static organization, fUSI reproduces hallmark resting state fMRI dynamics, including dominant anti-correlated coactivation patterns (CAPs) and a structured transition architecture that converges onto three stable attractor modes. Together, these results establish transcranial fUSI as a portable and scalable complement to fMRI for connectome-scale mapping of mouse brain networks.

## Introduction

Resting-state functional magnetic resonance imaging (rsfMRI) has been widely used for whole-brain, noninvasive mapping of the human functional connectome ^1^. Relying on the observation that spontaneous fMRI fluctuations form coherent resting-state networks (RSNs) ^2^, rsfMRI has shed light on the intrinsic organization of large-scale intrinsic brain activity ^3^, its temporal dynamics ^4^, and its disruption in neurological and psychiatric disorders ^5^. Notably, converging evidence suggests that functional connectome organization is evolutionarily conserved ^6, 7^. This conservation extends beyond functional topography to encompass structure-function relationships ^8^ and the dynamical architecture of RSNs ^9^. This evolutionary continuity has enabled mechanistic interrogation of human imaging findings in physiologically-accessible species such as rodents, establishing rsfMRI as a powerful tool for uncovering the physiological mechanisms underlying rsfMRI connectivity, and its alterations in brain disorders ^10, 11^. However, despite its translational potential, fMRI-based functional connectomics in rodents remains constrained by the high cost and fixed infrastructure of MRI, and by key practical limitations related to the scanner environment that restrict experimental flexibility and multimodal integration ^12^. As a result, scalable brain-wide functional connectome mapping in rodents remains largely limited to specialized MRI facilities.

Functional ultrasound imaging (fUSI) is a rapidly emerging hemodynamic neuroimaging modality that could help overcome these barriers ^13^. By measuring power Doppler fluctuations proportional to cerebral blood volume (CBV), fUSI provides a hemodynamic readout that is related to, but distinct from, BOLD contrast as measured with fMRI ^14^. Importantly, fUSI combines high spatiotemporal resolution (∼100 µm, 0.1 s) ^15, 16^ with a light, portable, near-silent acquisition setup ^17^, that may enable functional neuroimaging in highly versatile experimental setups. However, most fUSI applications in rodents have focused on evoked responses or sensory processing ^15, 18^, with intrinsic functional network mapping being limited to a few reports, typically relying on single-slice implementations ^19, 20^. As a result, robust demonstrations that transcranial fUSI can map brain-wide RSN in the mouse brain are lacking.

Here we test whether non-invasive transcranial fUSI enables quantitative, connectome-scale mapping of mouse RSNs. We then benchmark fUSI-derived RSNs against rsfMRI across experimental conditions and preprocessing pipelines. We show that fUSI robustly reproduces brainwide rsfMRI networks with strong cross-modal agreement, and clear anchoring to axonal anatomy along dominant structure-function axes. We also find that fUSI-derived networks exhibit structured dynamics characterized by a low-dimensional transition architecture converging onto a small set of coactivation-modes. These findings position transcranial fUSI as a versatile platform for scalable connectome-scale mapping in rodents.

## Results

### fUSI reveals brain-wide functional connectivity networks in the mouse brain

To determine whether fUSI can map large-scale RSNs in the mouse brain, we optimized a transcranial multi-slice acquisition protocol in wild-type C57 mice and developed a preprocessing pipeline mirroring established fMRI analytical frameworks (Fig. 1a-c). Our protocol uses a linear probe mounted on a motorized translation stage to sequentially sample four coronal planes (TR = 2.4 s), yielding an interleaved, non-contiguous volumetric acquisition with brain-wide coverage (Fig. 1b).

**Figure 1.**
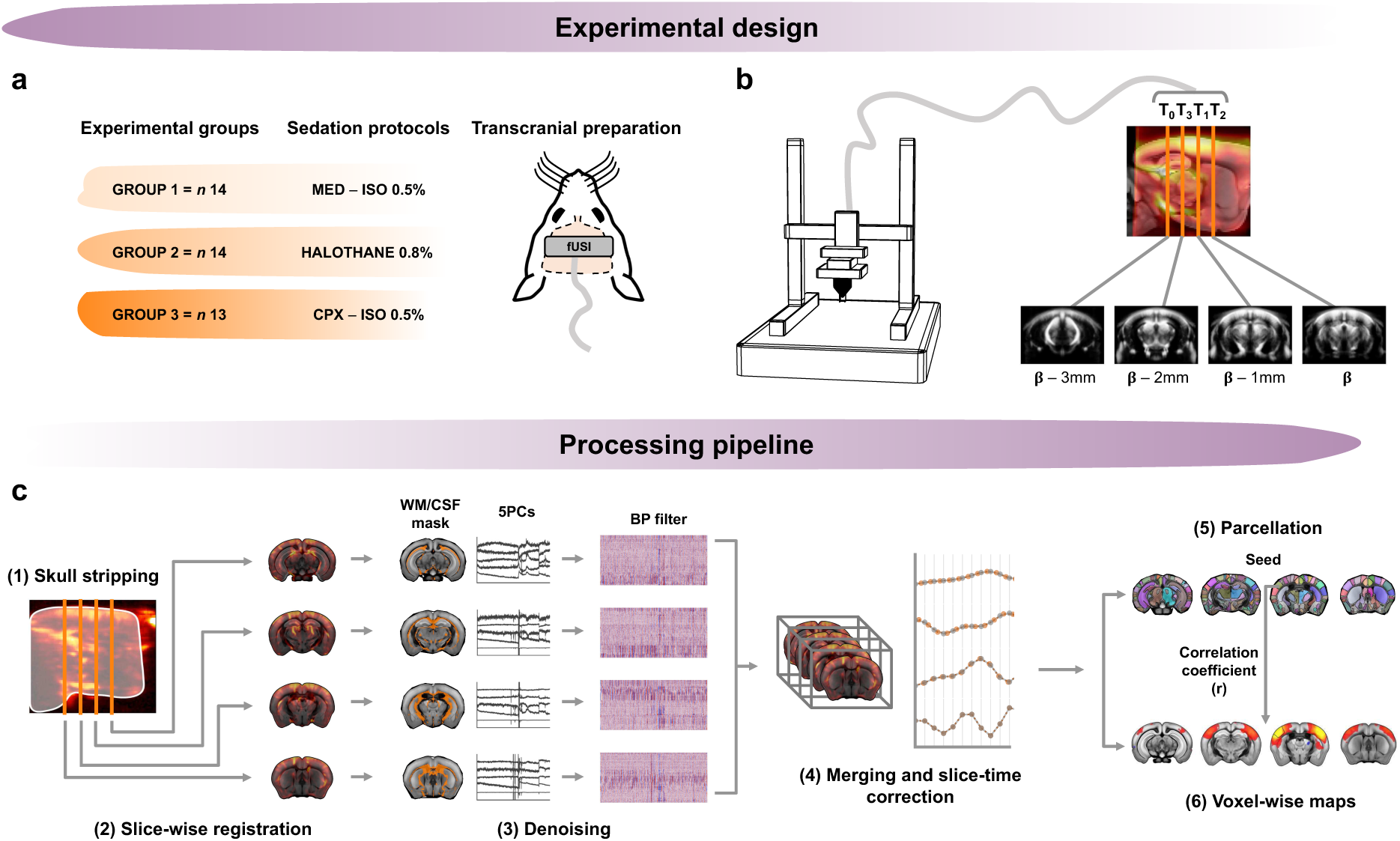
Experimental framework and analysis pipeline for resting-state fUSI network mapping. **(a)** Experimental design and transcranial preparation. Resting-state fUSI connectivity was acquired in three mouse groups under distinct light-sedation protocols: medetomidine-isoflurane (Med-Iso, n = 14), halothane (n = 15), and chlorprothixene-isoflurane (Cpx-Iso, n = 13), through the intact skull and skin. **(b)** Motorized multi-slice fUSI acquisition setup and imaging scheme. Brain coverage was obtained using four non-contiguous imaging planes positioned at −3, −2, −1, and 0 mm relative to bregma. Rightmost panel illustrates population average power Doppler images with corresponding slice locations. **(c)** Overview of the fUSI preprocessing pipeline. Data were processed slice-wise (1-2), denoised using anatomical CompCor (aCompCor) regressing the first five principal components derived from a white matter/cerebrospinal fluid (WM/CSF) mask, and band-pass filtered (3). Processed slices were then recombined into 3D volumes and slice-time corrected (4) prior to functional connectivity analysis (5-6).

As connectome-scale RSN mapping with fMRI is typically carried out under light sedation ^12, 21^, we performed fUSI mapping under three light-sedation regimens (Fig. 1a): (i) medetomidine combined with low-dose isoflurane (Med-Iso), a sedation mixture widely used in rodent fMRI ^22^; (ii) halothane, an inhalational anesthetic which at low doses preserves network organization in the rodent brain ^6^; and (iii) low-dose isoflurane + chlorprothixene microdosing (Cpx-Iso), a sedation protocol previously used in wide-field calcium imaging and large-scale electrophysiology ^23^, but not evaluated for brain-wide functional network mapping with fUSI.

To analyze resting-state fUSI networks, we developed an fMRI-inspired processing pipeline (Fig. 1c), encompassing slice-wise processing and nuisance regression tailored to non-contiguous slice sampling, band-pass filtering, volume reconstruction with slice-timing correction, and functional connectivity estimation. We then compared RSN detectability across sedation regimens using seed-based mapping.

This analysis revealed a set of robust, anatomically organized RSNs, recapitulating brain-wide network systems previously described in mice (Fig. 2 and Fig. S1a). Specifically, across regimens, fUSI consistently revealed a default-mode-like network (DMN), a laterocortical network (LCN), as well as robust thalamo-cortical coupling (Fig. S1, and Fig. 2a). Additional networks were also reliably detected, including hippocampal and subcortical systems (Fig. S1a). While RSN topography was broadly comparable across sedation regimens, the strength of functional connectivity varied substantially across conditions (Fig. S1a-b). Specifically, Cpx-Iso consistently revealed the strongest interhemispheric connectivity in somatosensory, motor, and thalamic regions (Fig. S1b, p<0.01, all comparisons, two-way repeated-measures ANOVA, FWER corrected) and the strongest anteroposterior connectivity within the LCN (Fig. S1b, p<0.001), whereas halothane and Med-Iso produced comparably weaker and/or more variable connectivity patterns across subjects (Fig. S1b). Cpx-Iso also showed evidence of thalamo-cortical and DMN-LCN anticorrelation. Based on these observations, we selected Cpx-Iso dataset for subsequent analyses (Fig. 2). Interestingly, in contrast to the field-standard Med-Iso regimen, Cpx-Iso (as well as halothane) also retained measurable spontaneous arousal-related fluctuations as assessed by pupillometry (Fig. S2), suggesting that these regimens preserve an arousal-linked dynamical component that is largely suppressed by medetomidine.

**Figure 2.**
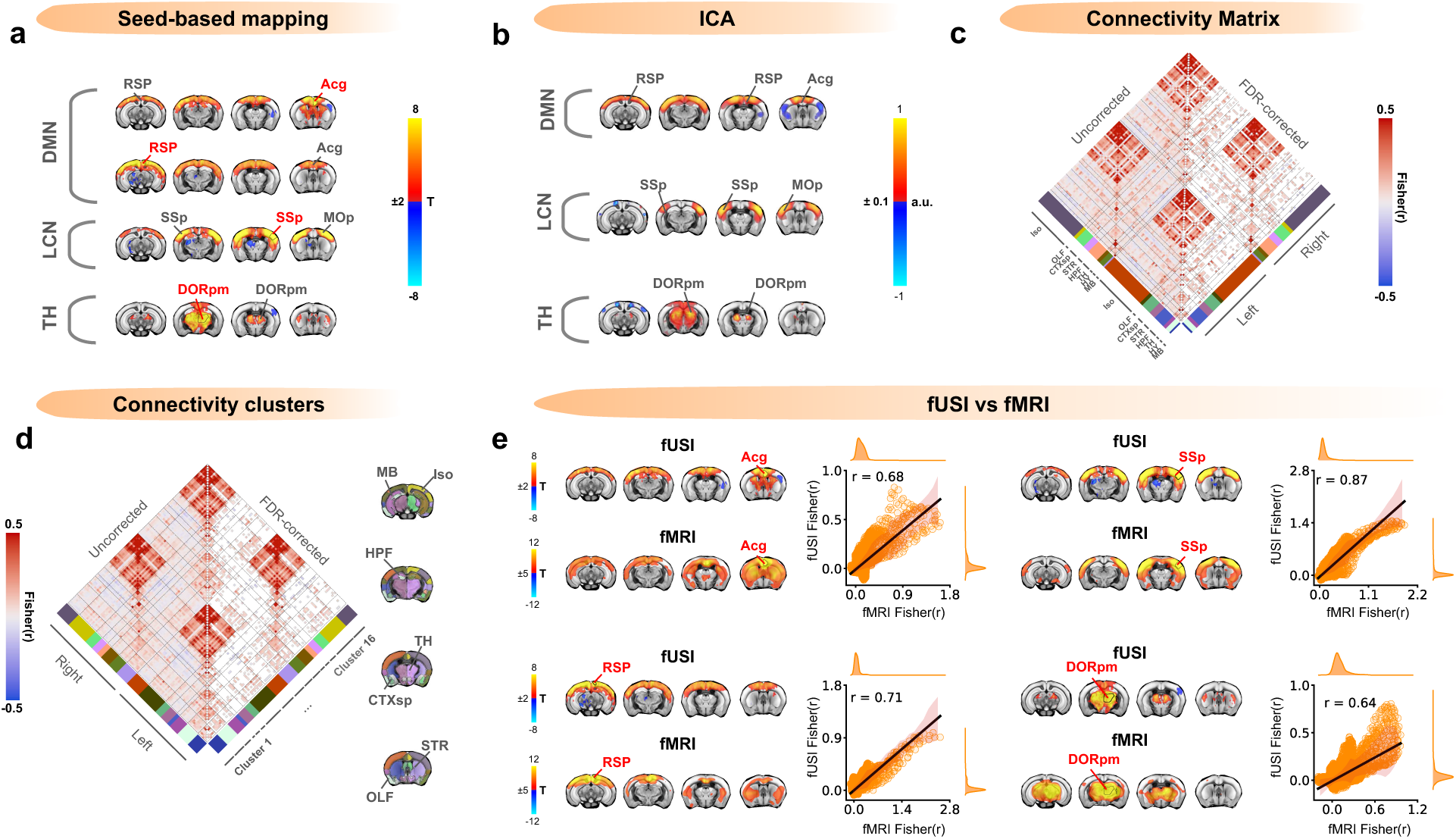
Transcranial fUSI maps distributed resting-state networks in the mouse brain. **(a)** Seed-based connectivity maps from unilateral seeds belonging to previously described RSNs (DMN, LCN, TH). **(b)** Group-level independent component analysis (ICA; k = 3) identifies a DMN, a LCN, and thalamic network. Maps are thresholded at 10% of the absolute component maximum. **(c)** Parcel-wise functional connectivity across 108 unilateral ROIs shown as group-average correlations (Fisher z; color scale). ROIs are assigned to eight macroregions (Iso, OLF, CTXsp, STR, HPF, TH, HY, and MB) based on Allen brain ontology and ordered by hemisphere. **(d)** Connectivity-based clustering of (c) yields 16 unilateral functional clusters, recapitulating the a priori parcellation in (c). Assignments are shown on the matrix and projected onto the Allen Brain Template. **(e)** Cross-modal seed-based comparison of fUSI- and fMRI-derived networks. Scatter plots report voxel-wise spatial correspondence between unthresholded Fisher-transformed seed maps. Maps are cluster-corrected, α = 0.05. Seed locations are indicated in red text labels. In (c-d), significance-masked connections are shown in the lower triangle (Wilcoxon test, Benjamini–Hochberg correction, q < 0.05). Acg, anterior cingulate cortex; RSP, retrosplenial cortex; SSp, primary somatosensory cortex; DORpm, polymodal thalamus; Iso, isocortex; OLF, olfactory areas; CTXsp, cortical subplate; STR, striatum; HPF, hippocampal formation; TH, thalamus; HY, hypothalamus; MB, midbrain.

Because preprocessing pipelines and nuisance regression are known to influence the correlation distribution and inferred functional organization of brain networks ^24, 25^, we systematically assessed how different nuisance regression strategies affect fUSI connectivity estimates in the selected Cpx-Iso dataset (Fig. S3). To this end, we compared the connectivity profile of primary somatosensory cortex (SSp) across ten nuisance-regression strategies (Fig. S3a; see Methods). fUSI connectivity patterns were largely consistent across denoising strategies, with the expected homotopic cortical topology remaining stable even under minimal or no nuisance regression (Fig. S3a). Among the tested approaches, anatomical CompCor (aCompCor) best preserved homotopic cortico-cortical connectivity while minimizing regression-induced anticorrelations (Fig. S3a,b) ^24^. This denoising approach also reduced between-subject variability in summary connectivity measures (Fig. S3c). We therefore adopted aCompCor as the reference denoising strategy throughout this work.

We next examined the architecture of fUSI-derived RSNs using complementary analyses (Fig. 2b–d). Independent component analysis yielded components highly consistent with seed-based mapping (Fig. 2a), revealing three major spatially coherent patterns corresponding to the DMN, LCN, and thalamic network (Fig. 2b). Notably, the DMN component showed opposing weights in midline versus lateral cortical territories, mirroring the DMN-LCN antagonism observed in seed-based maps and in rsfMRI. Connectivity matrices further support this organization, showing strong interhemispheric coupling (Fig. 2c), and a modular structure at the parcel level reconstituting known cortical and subcortical anatomical systems of the mouse brain (Fig. 2d). Collectively, these analyses show that transcranial fUSI robustly resolves established large-scale RSNs in the mouse brain.

### fUSI reproduces canonical large-scale rsfMRI networks

Having established that transcranial fUSI maps distributed functional networks in the resting mouse brain, we next examined how closely fUSI-derived networks recapitulate canonical fMRI systems. To this end, we leveraged an fMRI dataset (n=29) acquired under matched Cpx-Iso conditions as a benchmark for probing cross-modal correspondence. Seed-based connectivity mapping revealed strong spatial concordance between fUSI- and fMRI-derived DMN, LCN and thalamic networks (Fig. 2e). These networks also exhibited broadly comparable anatomical distribution in the two modalities, and robust voxel-wise inter-modality agreement (Acg, r=0.68; RSP, r=0.71; SSp, r=0.87; DORpm, r=0.64). Extending this analysis to additional cortical and subcortical regions, fUSI consistently recapitulated multiple fMRI network systems (including amygdala, hippocampal, striatal, insular networks), yielding similarly robust spatial correlations (Fig. S4; r = 0.62–0.76). Notably, these correspondences remained robust also after matching the fMRI sample size to the fUSI cohort (n=13 per modality; “sample-matched”; Fig. S5a, r=0.64-0.87), and after reprocessing rsfMRI data using an aCompCor-based denoising strategy matched to the fUSI pipeline (n=13; “pipeline-matched”; Fig. S5b). Under this harmonized preprocessing, fMRI networks displayed a modest reduction in spatial extent, yet cross-modal correspondence remained robust across all probed systems (r = 0.65–0.86, Fig. S5b). Taken together, these results show that transcranial fUSI recapitulates the large-scale architecture of the mouse functional connectome as probed with rsfMRI.

Interestingly, across matched conditions, fUSI yielded stronger estimates of cortico-cortical coupling, and interhemispheric (homotopic) connectivity than fMRI, most prominently within the laterocortical network (Fig S6a, p<0.001, FWER-corrected). In contrast, prefrontal-thalamic coupling (Acg-TH) was comparable across modalities, suggesting that this enhancement may not extend to cortico-subcortical links (Fig. S6a). Consistent with this, subject-level seed-based mapping showed more consistent cortical network detection in fUSI, with greatly reduced inter-subject dispersion of voxel-wise seed correlations relative to fMRI (Fig. S6b; KS tests p<0.001). However, fMRI exhibited broader spatial extent in subcortically distributed networks (Fig. S6b), suggesting a possible deeper brain coverage afforded by MR contrast. This pattern was also noticeable in our seed-based comparisons (i.e., Fig. 2e, Fig. S4), where fMRI revealed more spatially distributed subcortical network topographies than fUSI.

At the connectome level, cross-modal FC differences recapitulated this dissociation: fUSI showed markedly stronger cortico-cortical coupling and homotopic thalamic connectivity, whereas fMRI showed higher cortico-subcortical coupling, and higher homotopic connectivity in deeper brain regions (i.e., olfactory cortex, striatum, Fig. S7a). This profile is consistent with depth-dependent transcranial ultrasound attenuation (Fig. S7b), and with modality-specific contrast signal-to-noise ratio (SNR) characteristics that may differentially shape sensitivity across cortical versus deeper structures (Fig. S7b). Taken together, our cross-modal comparisons show that transcranial fUSI can detect canonical connectome-scale network systems in the mouse, with differential sensitivity to cortical versus deep subcortical organization compared to fMRI.

### fUSI networks are anchored to the structural connectome

fMRI connectivity studies have shown that mouse brain networks are closely tied to the underlying anatomical scaffold ^8^. We thus asked whether fUSI-derived networks show a comparable relationship to the underlying anatomical organization. We first quantified structure-function correspondence (SC-FC) by correlating the axonal structural connectome of the mouse brain with fUSI-derived functional connectivity (Fig. 3). Across regions, structural and functional connectivity showed a significant positive association (r=0.42; p<0.001, Fig. 3b). This value is in line with prior reports ^8^, and was only modestly lower than the corresponding coupling observed in fMRI data acquired under the same sedation conditions (Fig. S8a, r=0.51, p<0.001, fUSI vs fMRI interaction, p < 0.001).

**Figure 3.**
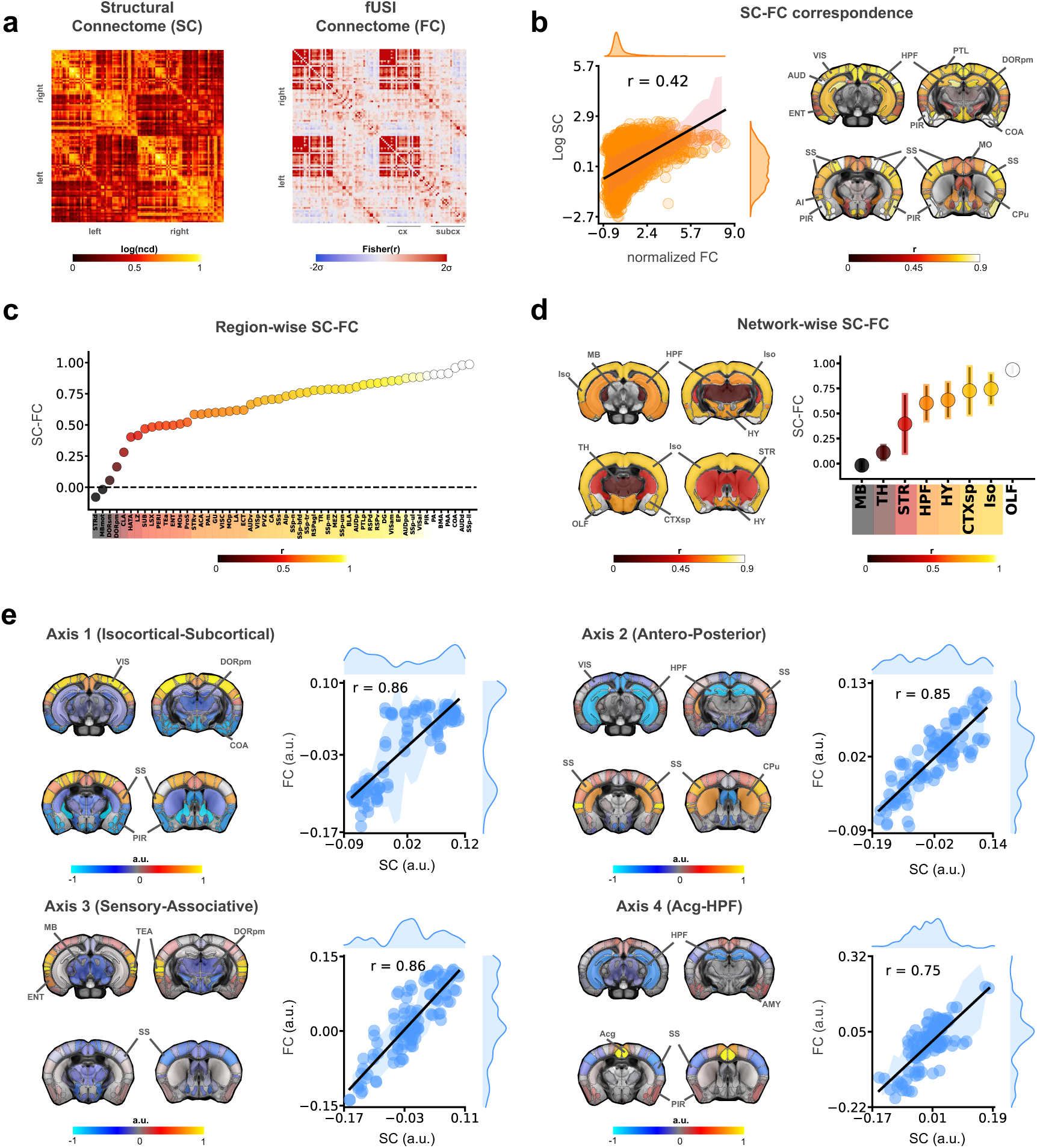
fUSI RSNs reveal multi-axis structure-function organization in the mouse connectome. **(a)** Structural (SC) and functional (FC) connectivity matrices across 108 unilateral ROIs. SC is shown as log-transformed normalized connectivity density (log(ncd)). FC is shown as Fisher z-transformed Pearson correlations. **(b)** Structure-function correspondence (SC-FC). Left, global SC-FC computed as the correlation between the lower triangles of SC and FC (shaded band, SD; marginals, univariate distributions of SC and FC edge weights). Right, ROI-wise SC-FC obtained by correlating the SC and FC profile of each ROI, and projecting the resulting values onto the Allen template. **(c)** Region-wise SC-FC derived from **(b)** for 54 bilateral ROIs (left/right averaged), ordered by correlation magnitude. (d) Network-wise SC-FC obtained by aggregating region-wise values into major macroregions (i.e., Iso, OLF, CTXsp, STR, HPF, TH, HY and MB). Brain map shows mean SC-FC correlation within each network. Summary plot reports corresponding mean SC-FC correlation values (mean ±SD). **(e)** Joint structural-functional axes from PCA on concatenated, normalized SC and FC matrices (Methods). Four components with strong SC-FC agreement are shown. Maps report ROI loadings, and scatter plots relate SC and FC loadings (line, fit; shaded band, SD). Abbreviations follow Allen nomenclature (Supplementary Table 1).

Brain-wide structure-function correlation, however, compresses spatially heterogeneous coupling into a single summary statistic, obscuring where, and in which systems, structure most strongly constrains function. We therefore computed region-wise SC-FC coupling by correlating for each brain region, its structural connectivity profile with its functional connectivity profile (Fig. 3b). We found that fUSI SC-FC correspondence was highly heterogeneous, with strongest coupling in olfactory, somatomotor and limbic territories, and weaker coupling in deeper striatal, thalamic and midbrain systems (Fig. 3b, c). Aggregating regions into major anatomical systems revealed a graded hierarchy of coupling (Fig. 3d), with the weakest correspondence in midbrain and thalamic systems (MB, r = −0.02; TH, r = 0.11), intermediate coupling in striatum, hippocampal, and hypothalamic networks (STR, r = 0.40; HPF, r = 0.60; HY, r = 0.63), and the strongest correspondence in cortical and olfactory systems (CTXsp, r = 0.73; ISO, r = 0.74; OLF, r = 0.94). Notably, applying the same analyses to fMRI produced a closely matching regional coupling profile (r=0.78; Fig. S8c,d) and a largely preserved system-level ranking (Fig. S8e), indicating that the observed coupling hierarchy is not specific to fUSI. Consistent with this concordance, modality-difference maps revealed only modest, spatially structured deviations, favoring striato-midbrain coupling in fMRI and lateral cortical, hypothalamic, and amygdalar coupling in fUSI (Fig. S8f). Together, these findings suggest that fUSI captures a conserved structure-function scaffold that is expressed with strong regional specificity, and marginal modality-dependent reweighting.

### Four structural–functional axes organize the mouse functional connectome

Measures of SC-FC coupling quantify the magnitude of structure-function agreement, but they do not reveal the geometry of this relationship, i.e., the latent dimensions along which anatomical wiring constrains functional interactions across systems. Given the marked regional heterogeneity of SC-FC coupling in fUSI, we therefore asked whether structure and function share a low-dimensional set of organizational axes. To address this, we concatenated normalized SC and FC matrices into a joint structural-functional connectivity matrix and applied principal component analysis to extract orthogonal components capturing variance shared across connectomes (Fig. S9). This approach enables the identification of multiple, coexisting independent SC-FC axes, providing a compact description of connectome organization, and revealing how different functional systems align with anatomical substrates.

We identified six principal components (PC) explaining most of the variance, each exceeding chance levels (bootstrap N = 10000; Fig. S10). Four axes showed tight correspondence between SC and FC loadings (r>0.70) and were retained for interpretation (Fig. 3e; Methods), whereas the remaining components were excluded because they were SC-dominated or exhibited poor SC-FC agreement (Fig. S10; Methods). Importantly, these axes revealed distinct organizational principles of the mouse connectome, each contrasting different sets of regions along mouse brain anatomy (Fig. 3e). Axis 1 (Isocortical-Subcortical; PC1, r = 0.86) delineates a cortical-subcortical gradient opposing widespread cortical areas to thalamic and striatal territories. Axis 2 (Antero-Posterior; PC3, r = 0.85) differentiates rostral from caudal cortical and subcortical structures. Axis 3 (Sensory-Associative; PC4, r = 0.86) separates primary sensory cortices from associative thalamic territories. Finally, axis 4 (Acg-HPF; PC6, r = 0.75) captures a limbic gradient opposing cingulate/medial prefrontal regions to hippocampal areas. These axes reveal paired functional systems that share strong anatomical anchoring, but differ in their covariation patterns. More broadly, these results show that structure-function correspondence in fUSI is organized along a small set of dissociable axes that differentially anchor functional systems to anatomical wiring.

### Recurring co-activation states shape fUSI network dynamics

fMRI studies have shown that spontaneous activity in rodents is organized into recurring, brain-wide activation states that organize moment-to-moment network dynamics ^26, 27^. To test whether fUSI exhibits comparable dynamics, we applied the co-activation pattern (CAP) framework ^27^. This approach identifies recurring brain states by clustering individual imaging frames based on whole-brain spatial similarity. We found that six CAPs accounted for most of the variance in fUSI recordings (VE=73%, Fig. 4, Fig. S11). As previously observed in fMRI, fUSI-derived CAPs revealed opposing patterns of coactivation (CAP/antiCAP pairs) characterized by inverse fUSI signal polarity (Fig. 4a). Following recent fMRI work ^9^, we refer to these paired states as coactivation modes (C-modes). In fUSI, C-modes occurred with unequal frequency (Fig. 4b), with C-Mode 1 and C-Mode 2 being more prevalent than C-Mode-3 (p < 0.001, Wilcoxon tests, FDR-corrected). fUSI CAPs also exhibited significant phase-locking to global signal fluctuations (Rayleigh test, p < 0.001, Bonferroni-corrected; Fig. 4c), thus reconstituting a key dynamic feature of fMRI timeseries ^9^. Interestingly, CAP/antiCAP pairs occurred at opposite phases of global signal fluctuation, supporting their interpretation as opposing states within a single underlying dynamical C-mode. Finally, consistent with the different sensitivity profile of fUSI and fMRI (cf. Fig. S7b), cross-modal CAP matching revealed good-to-excellent spatial overlap for prominently cortical CAPs (CAPs 1–4; r = 0.47–0.70; Fig. S12), whereas CAPs dominated by subcortical territories showed much weaker correspondence (CAP5–6; r≈0.16; Fig. S12). Together, these results show that fUSI recapitulates canonical co-activation dynamics described with fMRI.

**Figure 4.**
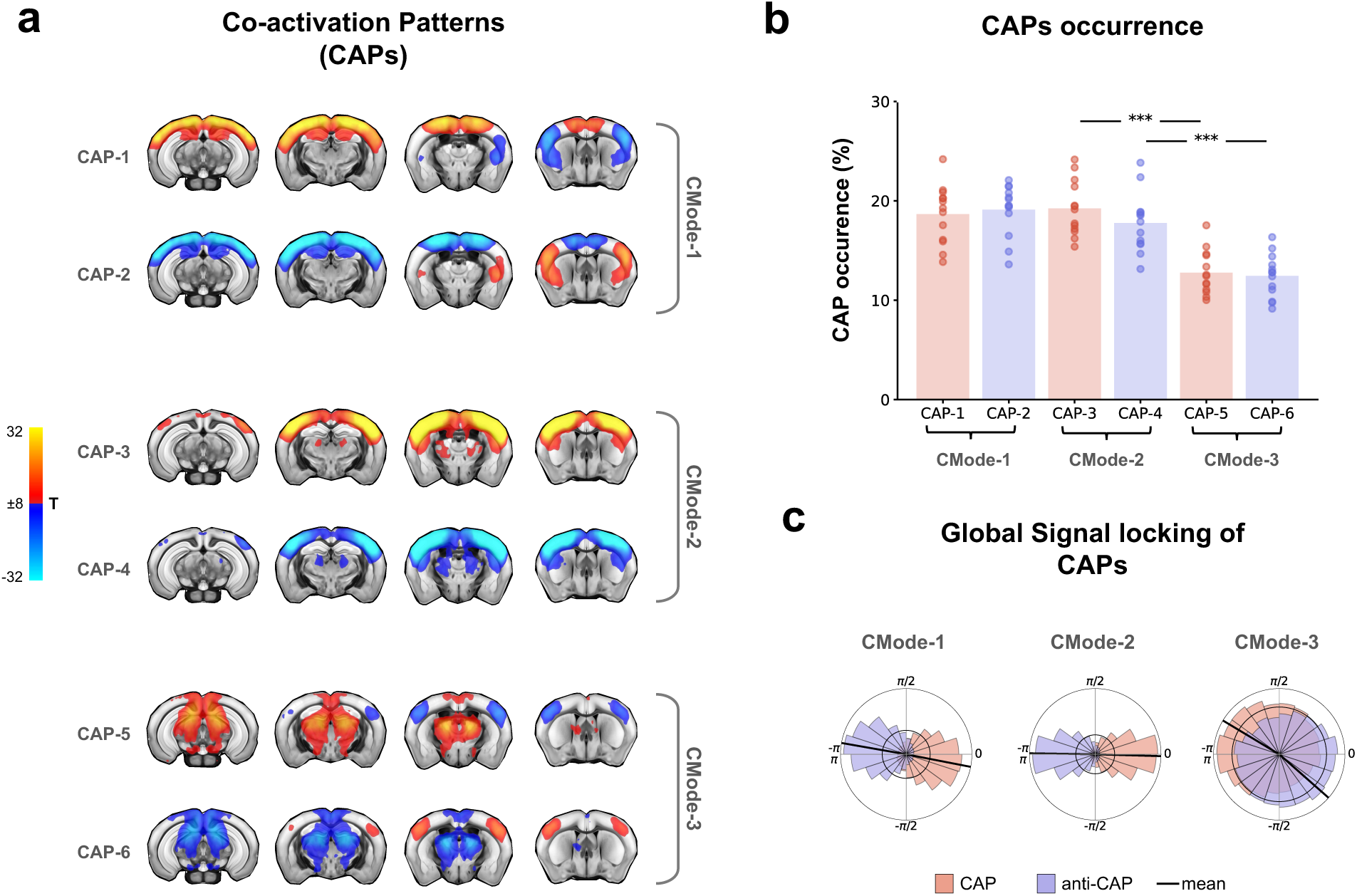
Recurring co-activation states structure fUSI network activity. **(a)** Co-activation patterns (CAPs) derived from resting-state fUSI (n = 13 mice) by framewise clustering (k = 6). The six CAPs form three oppositely signed CAP/antiCAP pairs (C-modes), expressed as group-level t-maps (warm colors, increased fUSI signal; cool colors, decreased signal; thresholded for display at |t| > 8, cluster-corrected). **(b)** CAP occurrence rates. Bars denote group mean, dots indicate individual mice. CAPs are grouped into C-modes; C-mode 1 and C-mode 2 occur more frequently than C-mode 3 (Wilcoxon tests; multiple-comparison corrected; ***p < 0.001). **(c)** Phase locking of CAPs occurrence to global signal fluctuations. Circular histograms show the phase distribution of CAP and antiCAP occurrences within each C-mode (black line, mean resultant vector). CAP/antiCAP pairs occur at approximately opposite phases, consistent with significant phase locking (Rayleigh test; multiple-comparison corrected; p < 0.001).

### CAP dynamics converge on three dominant attractor states

Prior fMRI work has shown that CAP dynamics is constrained by preferred transition paths quantified in a one-step transition space ^28^. This result raises the possibility that fUSI-derived network dynamics may similarly evolve through stereotyped trajectories. We therefore quantified CAP-to-CAP transitions after removing self-transitions to focus on state changes, and estimated the corresponding empirical transition probabilities using one-step transition matrices (Fig. 5a; Methods). While this analysis revealed multiple significantly preferred transitions, the inferred graph remained dense even under conservative thresholding (Fig. 5b; Fig. S13), limiting interpretability and obscuring higher-order dynamical motifs. Departing from our prior rsfMRI analyses, we thus examined a two-step transition model, which quantifies the probability of reaching a given CAP in two steps via any intermediate state (Fig. 5c). Strikingly, the resulting two-step transition landscape was markedly more structured. Specifically, across animals, trajectories preferentially linked each CAP to its opposing antiCAP, revealing three stable attractor-like motifs corresponding to the three C-modes defined above (Fig. 5c). Importantly, this organization was robust to different thresholding strategies (Fig. S13a). Moreover, application of this transition analysis to fMRI data recapitulated the same attractor structure (Fig. S13b), suggesting that this dynamic architecture is modality-invariant. Together, these findings provide a principled dynamical account for CAP/antiCAP pairing, consistent with the interpretation of each pair as opposing poles of a common fluctuating C-mode. More broadly, they also show that fUSI can reliably and accurately track connectome-scale RSN dynamics in the rodent brain.

**Figure 5.**
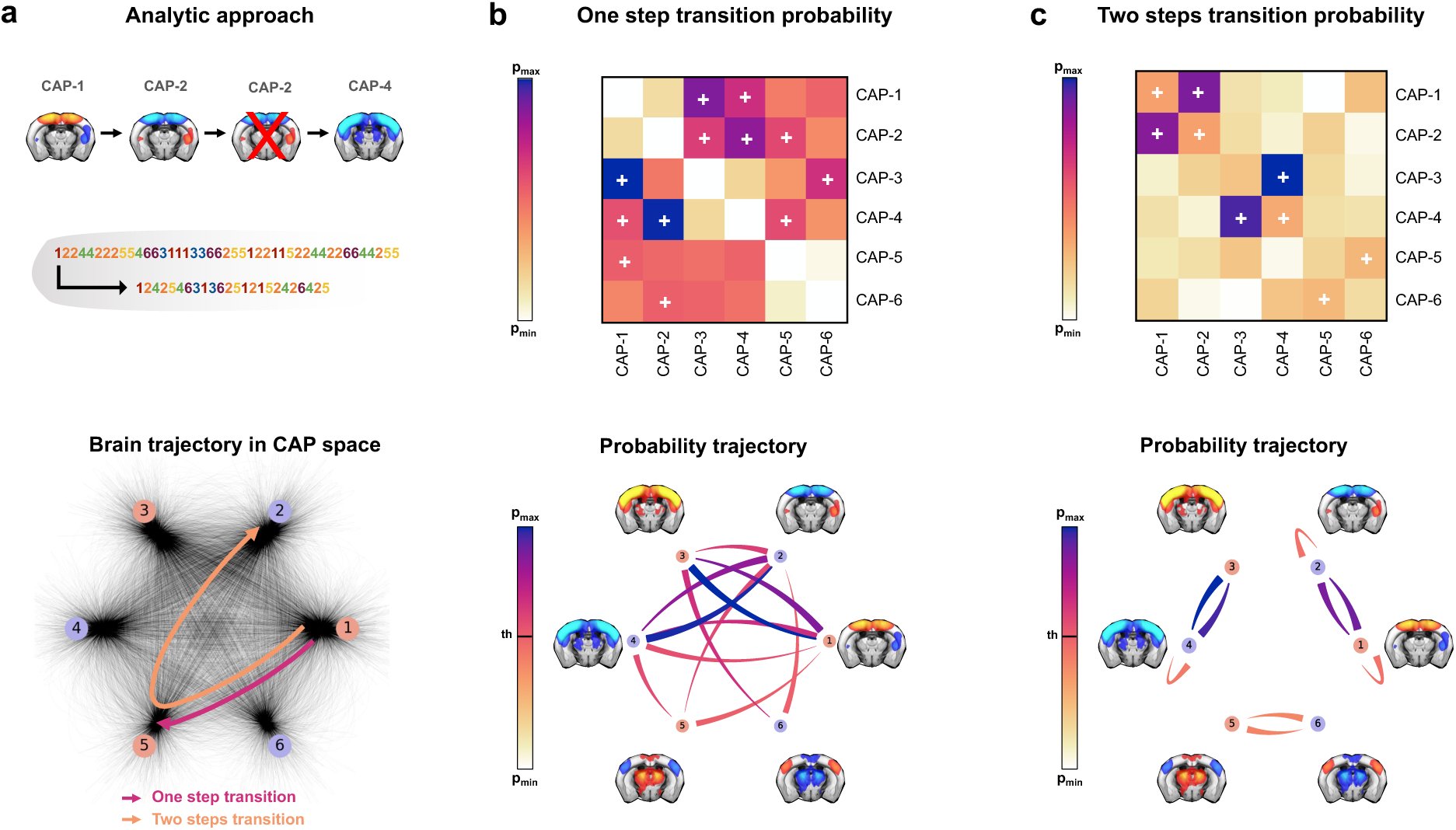
Transition architecture of coactivation patterns. **(a)** CAP transition analysis. Self-transitions were removed to isolate state changes (top), yielding a reduced CAP sequence used to compute one-step (magenta) and two-step (orange) transitions (bottom). **(b)** One-step transitions. Empirical one-step transition probability matrix (top) and corresponding directed graph after degree constrained (DC) thresholding (bottom). **(c)** Two-step transitions. Two-step transition probability matrix (top; probability of reaching a CAP in two steps via any intermediate state) and DC thresholded graph (bottom), highlighting preferential transitions between opposing CAP/antiCAP pairs. Plus symbols in matrices mark edges retained after DC thresholding (p<0.001). In bottom graphs in (b) and (c), nodes denote CAPs, and edge width and color are scaled proportionally to transition probability.

## Discussion

Here we report that multi-slice transcranial fUSI can reliably map the mouse functional connectome at brain-wide scale. We show that fUSI connectivity is anchored to the axonal connectome, and can be summarized by a set of joint structural-functional axes that capture dominant gradients of connectome organization. We also show that fUSI-derived networks show a highly structured transition architecture, converging upon three dominant coactivation modes. These findings highlight transcranial fUSI as a scalable neuroimaging modality for probing the intrinsic organization of functional brain networks in the mouse.

Previous work has shown the feasibility of using fUSI to map functional connectivity in rodents ^19, 20, 29^. However, most work to date has relied on single-plane acquisitions and restricted fields of view, limiting access to the distributed topology that defines brain-wide RSNs. Additional common constrains include the use of heterogeneous preparations and experimental settings, often involving non-resting conditions, or the use of invasive cranial windows, which can perturb baseline hemodynamics ^30, 31, 32, 33^. Together, these factors have hindered cross-species alignment of fUSI findings through comparative mapping of evolutionarily conserved, brain-wide network systems ^7, 11, 34^. By implementing fMRI-inspired sedation protocols and data preprocessing strategy, our work extends transcranial fUSI to connectome-scale mapping, revealing that this neuroimaging modality closely reproduces key organizational motifs that support RSN architecture across species. These include the presence of robust homotopic functional connectivity, as well as the presence of large-scale rodent precursor of the human networks, such default-mode-like network, and a latero-cortical system anticorrelated to the DMN (Fig. S3). Crucially, we also show that fUSI-derived networks exhibit dynamic organization recapitulating canonical features reported with fMRI ^9^. These findings expand the scope of fUSI as a platform for connectome-scale studies of RSN organization in rodents, complementary to fMRI.

While fUSI-fMRI correspondence was high across both cortical and subcortical networks, modality specific differences were also apparent. Specifically, fUSI provided considerably stronger and more reliable measures of cortico-cortical connectivity, resulting in more sensitive measures of network organization in these regions than rsfMRI. Consistent with this, fUSI-derived network patterns were evident even under minimal nuisance regression, indicating that connectome-scale fUSI structure in cortical regions is intrinsically robust. This feature may represent a practical advantage in light of the ongoing debate over the impact of specific preprocessing choices on fMRI-derived connectivity ^35^. By contrast, fMRI showed broader representation of deep cortical and subcortical regions, including striatal and hippocampal networks, and deep cortical areas, including insular components of the salience network ^6, 36^, for which fUSI did not identify a reliable correlate. This discrepancy is consistent with depth-dependent ultrasound signal attenuation ^37^, and may be exacerbated in brain regions such as the temporal/auditory cortex, or the striatum, where vessels are oriented approximately parallel to the ultrasound waveform ^13^. Notwithstanding these differences, the overall architecture of prominent subcortical networks (including hippocampal, striatal and midbrain), appeared to be consistent across modalities, suggesting that fUSI is well-suited to probe connectivity also in these regions. Further research will clarify whether skull removal, a preparation that improves deep-tissue sensitivity in fUSI ^38^, may yield better cross-modal matching, or whether these differences are contrast-dependent or modality specific.

A potential source of difference in the RSN organization mapped by fUSI with respect to fMRI, is the vascular nature of the hemodynamic signals these techniques are sensitive to. Theoretical and experimental work suggest that fUSI reflects a linear estimate of microvascular CBV fluctuations ^13^, while BOLD reflects a complex combination of vascular and neurometabolic contributors, including CBV, CBF and hemoglobin oxygenation ^39^. In this respect, predominantly CBV-weighted hemodynamic mapping may offer superior spatial specificity and greater fidelity to underlying neural activity. These benefits may reflect a closer relationship with local neuronal firing ^40^, together with reduced susceptibility to baseline-dependent metabolic effects that can decouple BOLD signal changes from oxygen metabolism ^41^. Moreover, CBV-weighted signals may be less affected by the spatial blurring introduced by large draining veins, a known limitation of conventional BOLD fMRI ^39^. fUSI could thus in principle permit relatively more specific localization of neuronal activity than BOLD fMRI. Although this added specificity did not confer an obvious advantage for mapping the widely distributed networks that support RSN activity in the mouse, it has already shown promise in fUSI-based mapping of sensory-evoked responses ^42^.

Beyond cross-modal benchmarking, our study also offers an updated view of how large-scale functional networks are shaped by anatomy, and organized around a low-dimensional set of dynamical states. At the connectome scale, global structural-functional coupling in fUSI closely matched prior mouse rsfMRI reports, corroborating the notion that RSNs are robustly constrained by the anatomical scaffold ^8, 43^. At the regional level, we observed a graded hierarchy with strong coupling in primary sensory and olfactory territories, and weaker coupling in associative, thalamic, and striatal regions. This organization is broadly consistent with macroscale unimodal-polymodal gradients reported in humans ^44, 45^ and rodents ^8^. Crucially, we extend classical cortico-centric graph embedding by extracting shared structural-functional axes and by showing that structure-function correspondence in the mouse brain is inherently multidimensional. Multiple, largely independent axes coexist, each capturing a distinct anatomical constraint (i.e., cortico-subcortical, antero-posterior, sensory-associative, and limbic). These findings extend gradient-based accounts to connectome-wide geometry, suggesting that rodent RSNs are organized along a small set of dissociable wiring motifs.

Importantly, beyond CAP topography, our work also resolved the transition structure between co-activation states. Specifically, by removing self-transitions to isolate genuine state changes, we uncovered a structured landscape in which two-step transitions preferentially connect each CAP to its antiCAP. Our observation that dynamic CAP transition converge onto three dominant attractor states supports recent conceptualizations of spatially-opposed CAPs as being constituents of a single coactivation-mode ^9^. Notably, the same transition architecture was recovered in matched fMRI data, indicating that this organization reflects a modality-invariant property of RSN dynamics. More broadly, our results are consistent with the emerging view that RSN dynamics can be parsimoniously explained by a small number of recurring spatiotemporal motifs ^46^. Our CAP-based framework offers an operational “transition-space” phenotype for quantifying, and comparing across modalities and species, the impact of perturbations on network dynamics using fUSI.

Reflecting prevailing practice in rodent rsfMRI connectomics ^8, 21^, here we performed fUSI RSN mapping using a light-sedation protocol. Pharmacological stabilization remains a widely utilized, pragmatic route to reproducible brain-wide network mapping, as it promotes stable and comparable brain states across animals and conditions ^12^. This is particularly important given accumulating evidence that awake head-fixed mice spontaneously traverse highly heterogeneous behavioral and arousal states that may diverge from the quiet-wakefulness typically sampled in human rsfMRI mapping ^47, 48^. Motivated by evidence of suboptimal network detection using medetomidine-isoflurane (a common rodent rsfMRI regimen), and the recent commercial discontinuation of halothane, we introduce a light sedation regimen (Cpx-Iso microdosing), that combines procedural simplicity with robust RSN mapping, while retaining measurable arousal-linked fluctuations.

Despite the practical advantages of sedation, there is a strong rationale for extending connectome-scale mapping to awake animals in both fMRI and fUSI ^12^. Accordingly, early demonstrations of brain-wide functional connectivity mapping in awake preparations have now been reported for both modalities ^20, 28, 31, 49^. However, in fUSI the transition from task- or sensory-evoked imaging to RSN mapping has proven more challenging than initially anticipated.

A primary limitation is the prevalence of large, transient motion-induced “flash-like” artefacts that contaminate awake recordings and strongly bias framewise RSN inferences ^50, 51^. Relative to what is typically encountered in rodent fMRI, these events appear more frequent, larger in amplitude, and more spatially pervasive, greatly increasing the burden on artefact detection and censoring ^28,50^. Moreover, transcranial acquisitions in awake animals are further constrained by Doppler contamination from extracranial and pericranial muscle activity, which reduces the usable lateral field of view, and greatly limits lateral coverage within coronal planes ^52^. Current efforts are therefore converging on combined solutions spanning physiology-aware denoising and artefact modelling ^31, 50^, ultra-stringent frame censoring, and in some settings, also acoustic optimizations that may require more invasive cranial access ^51, 53^, with initial encouraging results in small, highly curated datasets ^54^. Within this evolving landscape, our results provide a stable reference point for connectome-scale network topology and dynamics analyses of fUSI-derived networks, and a benchmark for method development toward awake fUSI connectomics.

In conclusion, our work demonstrates that transcranial fUSI can be used for connectome-scale mapping of intrinsic-network activity in the mouse. Beyond cross-modal validation with rsfMRI, our analyses show that these networks are differentially constrained by axonal wiring along a small set of shared structural-functional axes, and that their spontaneous activity is organized around recurring CAP states that converge onto three dominant coactivation modes. Our results provide a solid methodological foundation for reliable fUSI-derived RSN network mapping in rodents, positioning fUSI as an emerging platform for scalable rodent functional connectomics.

## Materials and Methods

### Ethical statement

Animal studies were conducted in accordance with the Italian Law (DL 26/2014, EU 63/2010, Ministero della Sanità, Roma) and the recommendations in the National Institutes of Health Guide for the care and use of laboratory animals. Animal research protocols were reviewed and consented to by the animal care committee of the Istituto Italiano di Tecnologia and the Italian Ministry of Health specifically approved the study protocol.

### Animals and experimental cohorts

All experiments were conducted in adult C57BL/6J mice (N = 50; Jackson Laboratory, strain 000664), including both males and females, aged 7–12 weeks. Animals were group-housed under controlled environmental conditions (temperature 21±1 °C; humidity 60±10%) on a 12 h light/dark cycle, with *ad libitum* access to food and water. To assess the robustness and generalizability of fUSI-derived functional networks across anesthetic regimens, experiments were performed in three independent cohorts using distinct light-sedation protocols. The first cohort (Med-Iso; males n=7, females n=8) received combined medetomidine-isoflurane anesthesia. Medetomidine was administered via continuous intravenous infusion (0.1 mg kg⁻¹ h⁻¹), following an initial tail-vein bolus (0.05 mg kg⁻¹), while isoflurane (0.5%) was delivered via endotracheal intubation, as previously described ^28^. The second cohort (Halothane; males n=10, females n=8) was anesthetized with halothane alone, administered via endotracheal intubation and maintained at 0.8%, following an established protocol ^55, 56^. The third cohort (Cpx-Iso; males n=9, females n=8) received a combination of chlorprothixene microdosing (Cpx) and low-dose isoflurane. Cpx (0.15 mg kg⁻¹; Sigma-Aldrich) was administered intramuscularly, and anesthesia was supplemented with 0.5% isoflurane delivered via a nose mask, as previously reported ^57, 58^. Cpx was prepared by dissolving the compound in saline (stock concentration 0.45 mg ml⁻¹) and freshly diluted on the day of injection to a final concentration of 0.045 mg ml⁻¹.

The 0.15 mg kg⁻¹ dose was applied uniformly across all animals in this study, and falls within a light-sedation range (0.15–0.3 mg kg⁻¹) established in preliminary experiments in adult mice with body weight ≥ 20 g. Cpx-Iso regimen has been previously used in wide-field calcium imaging ^23, 58, 59^ and electrophysiological studies ^60^ where it has been shown to stabilize neuronal activity without inducing burst suppression or compromising cortical responsiveness ^58, 61^.

### Resting-state fUSI

#### Animal preparation

Resting-state transcranial fUSI data were acquired in lightly sedated mice. Animal preparation procedures varied across experimental groups according to the anesthetic regimen (see below), while all other aspects of imaging preparation and data acquisition were kept constant. Across all protocols, an acoustic imaging window was prepared to optimize ultrasound transmission. The scalp was shaved, followed by brief application of depilatory cream (<1 min) to ensure complete fur removal, and the skin was thoroughly rinsed to prevent irritation ^62^.

*Group 1 (Med-Iso).* Mice were anesthetized with isoflurane (4% induction), intubated, and mechanically ventilated (2% during preparation). Animals were positioned in a stereotaxic frame for preparation of the imaging window and tail-vein cannulation. Medetomidine was administered as an initial intravenous bolus (0.05 mg kg⁻¹), followed after 5 min by continuous infusion (0.1 mg kg⁻¹ h⁻¹), while isoflurane was reduced to 0.5% for maintenance ^28^. Ventilation parameters were adjusted to maintain normophysiological arterial blood gases (paCO₂ < 40 mmHg; paO₂ > 90 mmHg, corresponding to >98% hemoglobin saturation).

*Group 2 (Halothane).* Mice were anesthetized with isoflurane (4% induction), intubated, and mechanically ventilated (2% during preparation). Following preparation of the imaging window, isoflurane was discontinued and replaced with halothane (0.8%). Ventilation parameters were matched to those used in Group 1.

*Group 3 (Cpx-Iso).* Mice were anesthetized with isoflurane (4% induction) and placed in a stereotaxic frame, after which isoflurane was reduced to 2% and delivered via nose cone. Following preparation of the imaging window, chlorprothixene (0.15 mg kg⁻¹, i.m.) was administered as a single bolus. Isoflurane concentration was then gradually decreased by 0.5% every 5 min to reach a final concentration of 0.5% after 15 min, which was maintained to ensure light sedation and preserved cerebrovascular autoregulation ^63^.

Across all groups, body temperature was maintained at 37 °C using a feedback-controlled heating pad. Heart rate and respiratory rate were continuously monitored (MouseOx, STARR Life Sciences). Pupil diameter was recorded concurrently to fUSI acquisition with a high-resolution camera (MRC Systems, 12M-i newSensor; see Pupillometry). Ultrasound gel (Aquasonic 100, Parker) was applied to the imaging window, and the ultrasonic probe was positioned approximately 1 mm above the skull using a four-axis motorized holder (Iconeus, Paris, France). Resting-state fUSI acquisition began 5 min after isoflurane concentration reached 0.5% in Groups 1 and 3, and 30 min after isoflurane discontinuation in Group 2.

### Ultrasound acquisition

Functional ultrasound imaging was performed using an Iconeus One scanner (Iconeus, Paris, France) equipped with a 15-MHz linear array probe (128 elements; 110 μm pitch; 8 mm elevation focal distance; 400 μm elevation focal width) mounted on a motorized translation stage^62^. Data were acquired at a pulse repetition frequency of 5,500 Hz using compounded plane-wave imaging across 11 tilted angles (−10° to +10° in 2° increments), yielding an effective sampling rate of 500 Hz after beamforming and compounding ^19, 20^. Power Doppler images were generated by applying spatiotemporal clutter filtering to stacks of 200 compounded frames. Tissue signal was removed using a singular value decomposition (SVD)–based filter that excluded the first 60 principal components ranked by variance) ^64^, resulting in Power Doppler images sampled at an effective rate of 2.5 Hz.

Multi-slice imaging was achieved by rapid motorized translation of the probe between slice positions (0.2 s per displacement). Combined with a 0.4 s Doppler integration time, this resulted in an effective interslice acquisition interval of 0.6 s. To ensure sufficient temporal resolution for resting-state analyses, imaging was restricted to four coronal slices, yielding one volume every 2.4s (temporal sampling rate ≈ 0.42 Hz). Each imaging session began with an angiographic scan covering the brain from β −5 mm to β +3 mm (81 slices; 0.1 mm interslice spacing; four repetitions per slice), producing a quasi-isotropic anatomical volume with a spatial resolution of 110 × 100 × 100 μm³. Probe positioning was verified using a vascular landmark defined by the bifurcation of the internal carotid artery into the anterior choroidal artery at β −1.8 mm ^31^. Resting-state acquisitions consisted of 60-min time series acquired using a four-slice protocol (β −3, −2, −1, and 0 mm), yielding 1,500 volumes with a spatial resolution of 110 × 100 × 400 μm³ and a repetition time of 2.4 s. Imaging sessions were randomized across experimental groups to minimize potential confounding effects.

### Preprocessing of fUSI timeseries

Resting-state fUSI data were preprocessed using a custom pipeline designed to parallel established fMRI workflows ^6^, while explicitly accounting for the spatial sparsity and temporal discontinuity inherent to multi-slice fUSI acquisitions. Two methodological features distinguish this pipeline: (1) a slice-wise preprocessing strategy in which each coronal slice is processed independently prior to volume reconstruction, and (2) systematic evaluation of nuisance regression strategies. Preprocessing was performed using AFNI ^65^, ANTs ^66^, FSL ^67^, and custom Python scripts (workflow summarized in Fig. 1c). Because multi-slice fUSI acquisitions consist of four non-contiguous coronal planes separated by 1 mm, functional data cannot be treated as a conventional 3D volume.

To ensure anatomical consistency while avoiding interpolation across missing tissue, we adopted a hybrid registration strategy combining 3D angiographic alignment with subsequent slice-wise functional registration. A custom angiographic fUSI template ^31^ was first manually aligned to the Allen Common Coordinate Framework (CCFv3), defining a common reference space (fUSI 100μm). The CCFv3-derived brain mask, parcellation, and white matter/cerebrospinal fluid (WM/CSF) masks were transformed into this space using the same registration parameters. Each subject’s angiographic scan was then registered in 3D to the fUSI 100μm template. Seven registration algorithms were evaluated for spatial normalization (ANTs Affine, ANTs SyN, ANTs Affine MINC, FSL FLIRT, FSL FNIRT, AFNI 3dAllineate, AFNI 3dQwarp), and the optimal transform was selected based on anatomical landmark correspondence. Across datasets, ANTs Affine (MINC implementation) consistently provided the most reliable alignment. The inverse transform of this alignment was subsequently applied to project the CCFv3 brain mask into each subject’s native angiographic space, enabling accurate skull stripping of the angiographic volume. Subject-specific four-slice templates were then generated by extracting the corresponding functional planes from each subject’s registered angiographic volume. Each slice was registered independently to its corresponding template using one of five algorithms (ANTs Affine, ANTs SyN, FSL FLIRT, AFNI 3dAllineate, AFNI 3dQwarp), selected on the best anatomical correspondence. The choice of a slice-wise preprocessing was motivated by two main considerations: (1) the large interslice gaps of our acquisitions precluded spatial interpolation between adjacent planes, and (2) motion-related fUSI artefacts consisting of occasional high-frequency flashes are largely independent across probe positions. All subsequent preprocessing steps were therefore performed independently for each coronal slice. Functional data were log-transformed to mitigate deformations caused by high-frequency intensity peaks characteristic of fUSI signals.

To identify an optimal nuisance regression strategy, ten alternative regressors were systematically evaluated across all three sedation datasets, spanning minimally aggressive, intermediate, and highly conservative approaches (Fig. S3). These included: no regression; mean signal regression from the lowest 5% intensity voxels (Low Intensity Signal); PCA-based regression using the five principal components obtained from non-brain tissue (Non-brain Tissue regression); mean signal regression from white matter (WM) and cerebrospinal fluid (CSF) (WM/CSF mask regression); global signal regression; PCA-based regression using the first principal component derived from all intracranial voxels (First mode regression); anatomical CompCor (aCompCor) ^68^, using the five principal components explaining the largest variance within a WM + CSF mask and temporal CompCor (tCompCor) using either the first principal component derived from all intracranial voxels (tCompCor N1P100), the first principal component from the most variable 2% of voxels (tCompCor N1P2), or the five principal components from the most variable 2% of voxels (tCompCor N5P2).

Qualitative benchmarking of this regression approach revealed three distinct classes of behavior (Fig. S3a). Minimally aggressive strategies (no regression and low-intensity signal regression) retained strong bilateral connectivity patterns but exhibited globally elevated correlation coefficients, consistent with residual nuisance variance. At the opposite extreme, aggressive strategies (global signal regression, first-mode regression, and most tCompCor variants) induced pronounced anticorrelations, man of which may by regression-induced, and as such devoid of physiological significance ^24^. Intermediate strategies (Non-brain Tissue, aCompCor, and tCompCor N5P2) yielded balanced correlation distributions without systematic shifts toward positive or negative values. Among these, aCompCor provided the most robust compromise between artefact suppression and preservation of biologically meaningful connectivity. Quantitative analyses supported these observations: voxel-wise correlation distributions for the primary somatosensory cortex (SSp) remained centered with limited tail expansion (Fig. S3b), and seed-to-seed connectivity values showed small between-subject variability relative to other strategies (Fig. S3c). Based on these observations and also recent fUSI work ^50^ we selected aCompCor as the default denoising strategy for our subsequent analyses.

Following regression, data were band-pass filtered (0.01–0.1 Hz; AFNI 3dBandpass). Pilot analyses carried out with a higher cutoff (0.3 Hz) did not reveal any qualitative improvement of our RSN mapping. To mitigate Gibbs artefacts and ensure comparability across sessions, the first five minutes and final three minutes of each recording were discarded, yielding 50-minute timeseries. Spatial smoothing was applied within each slice using a 2D Gaussian kernel (FWHM = 300 μm; AFNI 3dBlurInMask). Preprocessed slices were then reassembled into a sparse 3D volume using a custom slice-timing correction procedure based on linear interpolation to a common temporal reference. To further reduce the influence of transient artefacts, volumes were scrubbed if the global signal intensity in any slice exceeded three standard deviations from the median ^51^. On average, 10.98 ± 4.34 volumes (<1% of volumes) were removed per session. All recordings underwent comprehensive quality control, including assessment of the effectiveness of skull stripping, temporal signal-to-noise ratio, global signal correlation, frame stability, carpet plots, PCA scree plots, and spatial clustering patterns. A list of discarded scans and the corresponding reason for exclusion is reported in Supplementary Table 2.

### fUSI connectivity analysis

Resting-state functional connectivity in the mouse brain was computed in a set of 108 unilateral regions of interest (ROIs) derived from the Allen Brain Institute ontology (Supplementary Table 1). ROIs were selected to ensure broad anatomical coverage while maintaining sufficient spatial extent following resampling into the fUSI 100μm space (minimum extent ≥ 9 voxels). Voxelwise functional connectivity was first quantified using a seed-based mapping (SBM) approach. For each ROI, the mean seed timeseries was extracted from the corresponding seed voxels (FSL fslmeants) and correlated with the timeseries of all other brain voxels using Pearson’s correlation (AFNI 3dTcorr1D) ^65^. Subject-level correlation maps were Fisher z-transformed (AFNI 3dcalc), and group-level connectivity maps were estimated using one-sample t tests across individual z-maps (AFNI 3dttest++). Statistical significance was assessed with a voxelwise threshold of t > 2.1 followed by cluster correction (two-sided α = 0.05) implemented in FSL (cluster).

In complementary analyses, ROI-averaged time series were extracted for each animal and seed-to-seed functional connectivity was quantified using Pearson correlation between ROI time series, with correlation coefficients Fisher z-transformed prior to group-level analysis (Fig. S1b). Interhemispheric connectivity was quantified between homotopic ROIs, including primary somatosensory cortex (SSp left–right), primary motor cortex (MOp left–right), and polymodal thalamus (TH left–right), whereas anteroposterior connectivity was assessed between SSp–MOp and TH–SSp. Group differences were evaluated using two-way repeated-measures ANOVA with ROI pair and anesthetic condition as factors, followed by Tukey’s correction for multiple comparisons (GraphPad Prism). The same seed-to-seed connectivity framework was also applied to support the preprocessing pipeline comparison (Fig. S3c), focusing on interhemispheric SSp connectivity and thalamo-cortical connectivity (SSp-TH) across the ten nuisance regression strategies.

Group-level independent component analysis (ICA) was performed using the Dictionary Learning algorithm ^69^ implemented in Nilearn ^70^. ICA was run with default parameters and k = 3 components, chosen to recapitulate the three distributed resting-state networks identified by SBM. For visualization (Fig. 2b), independent components were scaled to their absolute maximum values and thresholded at 10% of this extremal value. Parcel-wise functional connectivity matrices were computed using the 108-ROI parcellation. Parcel timeseries were extracted using Nilearn masking tools, and pairwise Pearson correlations were computed (NumPy corrcoef) ^71^. Connectivity matrices were generated at the single-subject level, Fisher z-transformed, averaged across subjects, and expressed as correlation coefficients after inverse Fisher transformation for display. Significant edges were identified using mass-univariate Wilcoxon tests. P-value correction was implemented using the Benjamini–Hochberg procedure at q < 0.05. ROIs were ordered according to established a priori Allen ontology network membership ^72^. To identify reproducible network modules, the group-average parcel-wise connectivity matrix was clustered using Rastermap ^73^, applied in pattern-discovery mode following row-wise standardization. Rastermap applies sequentially dimensionality reduction (100 principal components) followed by regularized k-means clustering (k = 8). No lag window was used as the matrix lacks a temporal dimension. The number of clusters was selected to match the *a priori* network organization derived from Allen brain atlas ontology (Fig. 2c). Because Rastermap returns clusters spanning both hemispheres, each cluster was subsequently subdivided into left and right counterparts to follow Allen ontology conventions. This step enabled visualization of interhemispheric connectivity in the anti-diagonal block of the functional connectivity matrix.

### Pupillometry

Pupillometry was performed concurrently with fUSI acquisitions using a synchronized video recording setup. Pupil images were acquired with a high-resolution camera (MRC Systems, 12M-i newSensor with integrated LED illumination), synchronized to the fUSI sampling scheme to obtain one image per probe position. Pupil diameter was quantified offline using a custom semi-automated tracking algorithm. For each recording, a region of interest containing the pupil was manually defined. Within this region, the pupil center was identified as the minimum-intensity point, and pupil boundaries were estimated based on local intensity gradients along the horizontal and vertical axes. A single intensity threshold, defined relative to pupil and background intensities, was selected for each video to segment the pupil area. This threshold was held constant across frames, while pupil position and size were tracked iteratively over time. Frames were flagged as unreliable when segmentation failed to capture the pupil center or when abrupt, non-physiological changes in pupil area were detected; in such cases, the last valid estimate was propagated forward. Extracted pupil timeseries were visually inspected, and recordings that failed to meet predefined quality criteria were excluded: inclusion required stable pupil tracking over the full recording duration (3600 s), allowing for occasional brief segmentation failures that did not affect overall signal interpretation. Because pupil images were oversampled relative to the effective fUSI volume rate, pupillometry analyses were restricted to a single consistently reliable slice position (T1). The tracking algorithm performed robustly in the Cpx-Iso and halothane groups. In contrast, Med-Iso sedation markedly altered pupil dynamics, resulting in unstable pupil segmentation and reduced signal quality (Fig. S2); only eight recordings from this group met inclusion criteria. For visualization and quantitative analyses, we selected equal-sized subsets of high-quality, quality-matched recordings across groups (n = 8 per group). Pupil dynamics was quantified using three complementary measures: (1) the power spectral density of pupil diameter fluctuations, estimated using Welch’s method (SciPy) and normalized by total power, with the canonical resting-state frequency band (0.01–0.1 Hz) highlighted; (2) the median pupil diameter across the recording; and (3) signal complexity, quantified using approximate entropy ^74^. Across all metrics, Med-Iso recordings showed markedly reduced temporal variability, whereas halothane and Cpx-Iso conditions exhibited faster, more structured pupil fluctuations consistent with preserved arousal-related dynamics.

### fMRI acquisition

We performed rsfMRI acquisitions in N=29 adult male and female C57BL/6J mice aged 8-11 weeks. Animal preparation for rsfMRI scans was carried out under Cpx (0.15 mg/Kg intramuscular injection) and isoflurane (0.5% administered via intubation) light-sedation regime, as described above. Animal preparation was performed as previously described ^6, 27, 55^. rsfMRI image acquisition was performed on a 7 Tesla MRI scanner (Bruker, Ettlingen), using a 72 mm birdcage transmit coil and a 3-channel solenoid coil for signal reception. Paravision 6.01 software (Bruker, Ettlingen) was used to operate the scanner. BOLD rsfMRI timeseries (1920 volumes-32 min) were acquired using an echo-planar imaging (EPI) sequence with the following parameters: repetition time (TR) / time to echo (TE) 1000/15 ms, flip angle 60°, matrix 98 × 98, FOV 2.3 × 2.3 cm, 18 coronal slices, slice thickness 550 µm, bandwidth 250 KHz.

### fMRI preprocessing

fMRI preprocessing was performed following established pipelines previously described by our lab ^27, 55^. Briefly, the first 50 volumes of each timeseries were discarded to ensure signal stabilization. Data were subsequently despiked, motion corrected, skull stripped, and spatially registered to a standard fMRI EPI template. Motion parameters (three translations and three rotations) and the mean BOLD signal extracted from a ventricular mask were included as nuisance regressors and removed from the timeseries. Preprocessed data were band-pass filtered between 0.01 and 0.1 Hz and spatially smoothed using a Gaussian kernel with a full width at half maximum of 500 μm. Additional frame-wise scrubbing was applied using a framewise displacement threshold of 0.05 mm. To enable direct comparison between fMRI and fUSI connectivity, preprocessed fMRI volumes were resampled into the fUSI100 μm reference space using AFNI 3dAllineate and restricted to the four coronal planes corresponding to the fUSI imaging slices. These slices were subsequently recombined to generate a sparse four-slice fMRI volume, thereby matching the spatial structure of the fUSI data. Seed-based functional connectivity analysis was then performed on the fMRI dataset using the same ROIs and procedures described above for fUSI.

For cross-modal comparisons, we retained the full rsfMRI dataset (n = 29) to allow for a robust assessment of RNS organization under using a sample size representative of the experimental conditions employed (i.e., 7T, no cryoprobe). Because the full fMRI dataset comprised a larger number of animals than the fUSI dataset, we used arbitrarily thresholded fMRI T-maps at t = 5 to facilitate qualitative comparison of spatial topology. In subsequent analyses, we randomly subsampled the rsfMRI dataset to match the fUSI sample size (n=13 each). This enabled visualization of *t*-maps using comparable threshold and saturation settings (Fig. S5). Furthermore, because fUSI data were denoised using aCompCor, whereas standard rsfMRI preprocessing in the mouse does not typically encompass this denoising strategy ^22^, we reanalyzed our rsfMRI datasets using a fUSI-matched pipeline. In this matched pipeline, slices were preprocessed independently to mirror the fUSI workflow. Individual slices were subsequently recombined into a four-slice rsfMRI volume without slice-timing correction. Across analyses, three versions of the fMRI dataset were therefore used as controls: a standard dataset (n = 29) processed with the standard fMRI pipeline; a randomly subsampled dataset (n = 13) processed with the standard fMRI pipeline (sample-matched dataset); and a randomly subsampled, preprocessing-matched dataset (n = 13) processed using the fUSI-specific preprocessing pipeline (pipeline-matched dataset).

### fUSI-fMRI comparison

To assess the correspondence between RSNs derived from fUSI and fMRI, we performed voxel-wise comparisons of seed-based connectivity maps obtained from the two modalities. Analyses focused primarily on four anatomically defined seed regions used throughout the fUSI analyses: ACg, RSP, SSp, and DORpm. These regions were selected as representative hubs of canonical resting-state networks in the mouse brain ^6, 22^. The same voxel-wise comparison framework was extended to additional cortical and subcortical seed regions, to probe generalizability of the observed relationships (Fig. S4). Spatial correspondence between fUSI and fMRI connectivity patterns was quantified by computing voxel-wise Pearson correlation coefficients between unthresholded, Fisher z-transformed seed-based maps. Linear relationships between modalities were characterized using the *pearsonr* function from the NumPy package. The standard error of the fit was evaluated piecewise by dividing the data into 20 segments. Marginal distributions of connectivity values were visualized using Gaussian kernel density estimation and projected onto the corresponding axes. For the four representative seeds, cross-modal correspondence was further assessed using both the subsampled fMRI dataset and the pipeline-matched dataset (n = 13). In the subsampled dataset, spatial correlations between fUSI and fMRI maps were estimated using bootstrap resampling (10,000 iterations, 13 animals selected randomly with repetition), yielding stable population-level estimates of cross-modal similarity (ACg: r = 0.65, 95% CI [0.61, 0.71]; SSp: r = 0.86, 95% CI [0.83, 0.88]; RSP: r = 0.59, 95% CI [0.50, 0.66]; DORpm: r = 0.63, 95% CI [0.59, 0.66]). For visualization purposes, Fig. S5a displays a single representative realization of random sampling without replacement, illustrating that when sample size and statistical thresholds are matched, fUSI- and fMRI-derived connectivity maps exhibit comparable spatial topologies.

### Sensitivity analysis

To compare the sensitivity of fUSI and fMRI for resolving large-scale functional connectivity, we first performed a group-level analysis of pairwise connectivity strength across modalities. Seed-to-seed functional connectivity was compared between fUSI and a standard fMRI dataset, as well as between fUSI and a pipeline-matched fMRI dataset, in both interhemispheric homologous regions, as well as for intra-hemispheric areas, or for cortical-subcortical components of the DMN (Acg-TH). Group-level statistical comparisons were performed on Fisher-transformed correlation coefficients (z values) using multiple unpaired t tests with false discovery rate correction (Benjamini–Hochberg, q = 0.05) in GraphPad Prism.

To probe the capability of each modality to resolve functional connectivity at the single-subject level we computed frequency seed-based maps that quantified the percentage of animals expressing above-threshold connectivity at each voxel (Fig. S6b). This was achieved by computing, for each subject and seed, the modified z-score (a robust z-score suitable for skewed distributions) of voxelwise seed-based map values and thresholding the resulting maps at 2.5. This procedure yields binarized subject-level seed-based maps in which only voxels showing strong seed correlation, operationally defining the corresponding network, are retained. The choice of a modified z-score threshold follows common robust heuristics for identifying extreme values (typically ∼3, ^75^). We used a slightly more permissive threshold (2.5), which we selected empirically based on visual inspection of thresholded maps to achieve coherent, anatomically plausible network delineation across subjects. We then computed the percentage of subjects who exhibited above-threshold connectivity at each voxel. To quantify modality-dependent differences in the distribution of seed-based connectivity maps, we compared the voxelwise seed-based correlation coefficient distributions obtained from fUSI and fMRI for each probed seed. Distributions pooled across subjects were tested for significant differences between modalities using a two-sided Kolmogorov–Smirnov (KS) test.

To assess modality-specific effects at the network level, we computed differences between FC matrices derived from each modality. We next examined how individual network edges differed between fUSI and fMRI by computing homotopic (i.e., within network) connectivity for selected pairs of cortical or subcortical regions. Between-network connectivity was quantified unilaterally in the left hemisphere. For each network pair, we calculated the mean of all pairwise correlations contained in the rectangular block linking the two networks, corresponding to the average connectivity between every ROI in network A (left hemisphere) and every ROI in network B (left hemisphere).

Finally, we investigated whether these observed differences could be explained by variations in raw signal properties. To this end, we computed several metrics to characterize signal properties in both fUSI and fMRI (Fig. S7b). Images were analyzed after registration (merged without denoising for fUSI; resliced and split for fMRI), and for each voxel we calculated:

- Mean intensity defined as the average of a voxel’s time series calculated using only the 50% of time points that are below its median. Focusing on these lower-intensity points provides a conservative scrubbing approach to reduce bias from transient bursts, such as fUSI flash artifacts.
- Rayleigh contrast: 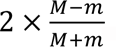, where *M* and *m* are the maximum and minimum values of the voxel time series, respectively.
- Temporal signal-to-noise ratio (tSNR), defined as the ratio of the mean to the standard deviation of the voxel time series.

We display voxelwise maps of all three metrics in Fig. S7. For each map, the corresponding value distributions are reported on the side. Because of fundamental differences in signal units between modalities, the distributions of average signal values were standardized to their respective minimum and maximum. Profiles of each metric along the vertical axis are also displayed, computed by averaging across the anteroposterior and mediolateral dimensions; these profiles are shown in black.

### Structural-functional connectivity (SC-FC) analysis

Structural-functional connectivity (SC-FC) analyses were performed by integrating mesoscale structural connectivity data from the Allen Mouse Brain Connectivity Atlas with functional connectivity derived from our fUSI dataset acquired under the Cpx-Iso sedation protocol. Structural connectivity (SC) estimates were based on a high-resolution (100 μm³) voxel-wise model of the mouse brain connectome ^76, 77^.

A SC matrix was constructed using the allensdk Python library based on the same parcellation scheme employed for functional analyses (Supplementary Table 1). For each ROI, first-order monosynaptic SC was quantified using normalized connectivity density (ncd). To facilitate direct comparison with functional connectivity (FC), which is intrinsically undirected, input and output structural connections were averaged to yield a symmetric SC matrix. The FC matrix was computed by calculating Pearson correlation coefficients between ROI-averaged time series for all pairs of ROIs in the 108-region parcellation. To ensure comparability between structural and functional measures, SC values were log-transformed, whereas FC values were Fisher z-transformed prior to analysis. As a first step toward quantifying structure-function coupling at the global level, we computed the brain-wide SC-FC correspondence as the Pearson correlation between the lower triangles of the SC and FC matrices. This procedure yielded a single scalar measure of global structure-function coupling (Fig. 3b). An analogous analysis was also performed on the fMRI dataset (n = 29).

To further investigate structure-function correspondence at a finer spatial scale, we quantified SC-FC coupling on a per-region basis. For each ROI, we computed the correlation between its functional connectivity profile (FC with the remaining 107 ROIs) and its structural connectivity profile (ncd with the remaining 107 ROIs). These region-wise SC-FC correlation values were mapped onto the atlas parcellation to visualize the spatial distribution of structure-function concordance across the brain. To assess how SC-FC coupling in distinct network systems, the resulting values were grouped and averaged into networks corresponding to the upper-level macroregions of the Allen Brain Institute ontology. A corresponding analysis was also performed on the fMRI dataset. Region-wise SC-FC values (averaged across left and right homologous ROIs) were also Pearson-correlated across modalities, to probe cross-modal correspondence between the two methods. To test whether the relationship between SC and FC differed between fUSI and fMRI, we also fitted a linear model including SC, imaging modality, and their interaction (FC ∼ SC × modality), equivalent to an analysis of covariance (ANCOVA). The interaction term revealed a significantly lower SC-FC slope for fUSI compared to fMRI (Δslope = −0.094, p < 0.001).

### Structural-functional axes

Motivated by the assumption that structure-function correspondence is not uniform across the brain, we sought to identify subnetworks in which anatomical and functional connectivity were differentially coupled using a multivariate approach. To this end, structural (SC) and functional (FC) connectivity matrices were concatenated along the row dimension after mean-centering and variance scaling, yielding a joint structural-functional connectivity matrix (SC-FC matrix). In this representation, each row corresponds to a single ROI and encodes its complete connectivity profile across both structural and functional domains relative to all other regions. To uncover dominant axes of shared structure-function organization, PCA was applied to the SC-FC matrix using singular value decomposition, after row-wise centering. This dimensionality reduction identifies orthogonal components that capture shared variance in SC and FC profiles across ROIs. Each principal component is defined by a left singular vector, reflecting the contribution of individual ROIs to the component, a right singular vector describing the corresponding connectivity pattern, and a singular value quantifying the variance explained. Owing to the concatenated structure of the SC-FC matrix, the right singular vector of each component can be partitioned into structural and functional segments. The degree of structure-function alignment along each axis was quantified by computing the Pearson correlation between these structural and functional segments, thereby identifying components exhibiting strong structure-function correspondence. The statistical relevance of the extracted components was assessed using a bootstrap-based null model. The SC-FC matrix was randomly permuted 10,000 times, and PCA was applied to each shuffled matrix to generate a null distribution of explained variance. Figure S9 shows the average null scree plot (orange) together with the 95th percentile of the null distribution (blue). Components whose explained variance exceeded this threshold were considered to capture structure-function covariance above chance level. Using this criterion, six components were identified as significant.

Figure S10 reports the loadings of these six components together with the correlation between the structural and functional segments of their right singular vectors. Four components exhibited robust structure-function correspondence (r>0.6), indicating axes along which anatomical and functional connectivity profiles are closely aligned. Component 2 was dominated by the structural connectome, and likely reflects the intrinsic left-right symmetry of the Allen Brain Institute structural connectivity data ^77^. In contrast, component 5 revealed a marked decoupling between structural and functional connectivity in subcortical regions, particularly within the midbrain and thalamus, where anatomical and functional profiles diverged substantially, consistent with patterns observed in the regional SC–FC analyses. This component was not included in further analyses.

### Coactivation patterns (CAPs)

To investigate the spatiotemporal organization of fUSI RSNs, we applied the co-activation patterns (CAPs) framework originally developed for rsfMRI timeseries ^27, 28^. This approach identifies recurring, brain-wide patterns of activity by clustering individual frames based on their spatial similarity. Following previously established procedures ^27^, all preprocessed fUSI timeseries from 13 animals were concatenated into a single Casorati matrix (n_voxels × n_timepoints). fUSI timeseries were preprocessed as described above, with the exception that nuisance regression was omitted in order to preserve the global component of hemodynamic fluctuations, which is known to play a critical role in CAP structure ^27^. Voxelwise timecourses were mean-centered and variance-normalized prior to clustering. CAPs were identified using k-means clustering implemented in MATLAB, with correlation distance and k-means++ initialization, run for 500 iterations ^27, 28^. To reduce sensitivity to local minima, the entire clustering procedure was repeated independently three times, and the solution explaining the largest proportion of variance was retained for subsequent analyses. For visualization of CAP spatial patterns, t-maps of normalized fUSI activation were computed across all frames assigned to each cluster. These maps were thresholded at T > 8 (p<0.001) and cluster-corrected using FSL (cluster). The optimal number of clusters was determined by systematically varying k from 2 to 18 and computing, for each value, the proportion of variance explained, defined as the ratio of between-cluster variance to total variance ^27^. Within-cluster variance was calculated as the average, across clusters, of the sum of squared distances between frames and their respective cluster centroids, whereas between-cluster variance was defined as the average squared distance between each cluster centroid and the global data centroid. Based on this analysis, k = 6 clusters were selected for three reasons. First, prior rsfMRI studies have shown that six clusters capture the majority of spontaneous brain dynamics across datasets, accounting for more than 75% of the variance ^27^. Second, the variance-explained curve for fUSI exhibited a clear elbow at k = 6, with higher-order clusters (k > 6) contributing less than 1% additional explained variance (Fig. S11). Third, selecting an even number of clusters preserves interpretable CAP–antiCAP pairs, referred to as C-modes, consistent with recent conceptualizations of CAPs ^9^.

For each subject, the occurrence rate of each CAP was computed as the proportion of frames assigned to that cluster. Group-level occurrence rates (Fig. 4b) were obtained by averaging across subjects, and differences in occurrence across CAPs were assessed using the Wilcoxon-Mann-Whitney U test, with multiple comparisons controlled using the Benjamini-Hochberg false discovery rate procedure. The temporal relationship between CAP expression and the global signal (GS) was further examined by analyzing CAP phase locking. For each CAP, a timeseries was obtained by correlating the CAP centroid with each frame of the concatenated timeseries. Instantaneous phase relationships were then estimated by computing the phase difference between the Hilbert transforms of the GS and the corresponding CAP timeseries. Phase differences were visualized as polar histograms with a fixed number of bins, alongside a matched uniform angular distribution. For each CAP, the mean phase difference was computed using the circular mean (circmean) from the scipy.stats module. Phase locking was formally assessed using the Rayleigh test, with Bonferroni correction for multiple comparisons.

### CAP dynamics

To characterize the dynamic interactions among the six identified CAPs, we developed a transition-based framework incorporating both one-step and two-step transition probability models. To emphasize meaningful state changes, all self-transitions (i.e., transitions from a CAP to itself) were excluded from both models. For example, a sequence such as 2-4-4-3-3-3-3-1-1-1-1-2-2-6-6-4-5-5-5-1 was reduced to 2-4-3-1-2-6-4-5-1 after removing repeated consecutive states. This transformation was motivated by the low-pass filtering of the data below 0.1 Hz, which makes self-transitions the most frequent events (accounting for approximately 54% of all transitions). Removing self-transitions allows the analysis to focus on transitions between distinct state changes, resulting in a reduced “transition space.” Operationally, this procedure alters the instantaneous sampling rate and facilitates the identification of pseudo-periodic transition patterns. Within this reduced transition space, we computed transition probability matrices. The one-step transition matrix was defined as the probability of direct transitions from CAP_i_ to CAP_j_, normalized by the total number of transitions. The two-step transition matrix captured the probability of transitioning from CAP_i_ to CAP_k_ through any intermediate CAP_j_. We note here that removal of self-transition results in i≠j and j≠k, but releases the constraint on i=k. Both transition matrices were visualized directly (Fig. 5, where (+) signs indicate edges above threshold, see below) and represented as directed graphs in which nodes correspond to CAPs and edges are weighted by transition probability, encoded through both edge width and color.

CAP transition matrices were thresholded using four different strategies, characterized by increasing stringency. Specifically, thresholds were set as (1) the 95^th^ percentile of a data-driven null distribution (th_95_); (2) the 99^th^ percentile of the null distribution (th_99_); (3) a chance-level threshold (th_Chance_); and (4) a degree constrained thresholding (th_DC_). The th_Chance_ was defined based on a uniform transition model, corresponding to the expected transition probability if all allowed transitions were equally likely. For the one-step transition matrix, self-transitions were excluded by construction, yielding th_Chance_ = 1/(n(n-1)), where n is the number of CAPs. For the two-step transition matrix, self-transitions were allowed, resulting in and th_Chance_ = 1/n^2. The th_DC_ corresponds to the maximal threshold that preserves at least one incoming and one outgoing edge for each node, thereby excluding sources and sinks and ensuring that all nodes participate in feedback-capable pathways.

Threshold levels were next determined using bootstrap-based null model. To this end, CAP state timeseries (i.e., the sequence of CAP assignments across volumes) were randomly permuted 10,000 times. For each permutation, self-transitions were removed and transition probability matrices were recomputed, generating null distributions for both one-step and two-step transitions. Figure S13a shows these null distributions, which consistently exhibited the ordering th_95_ < th_99_ < th_Chance_ < th_DC_. We report in Figure S13b the resulting subnetworks obtained under each thresholding strategy. Among the tested thresholds, th_DC_ preserved the most salient dynamical features of the CAP transition structure, including dominant cyclic motifs in the one-step transition matrix and attractor-like configurations in the two-step transition matrix, while avoiding excessive graph densification. While more permissive thresholding revealed additional structure, the convergence between CAP and anti-CAP organization remained dominant. Based on these considerations, we visualized transition graphs using th_DC._

### fMRI CAPs

To assess how modality-specific sensitivity influences CAP topographies, we additionally derived CAPs from the fMRI dataset (standard preprocessing, n = 29) using the same number of clusters (k = 6) as for the fUSI analyses. CAP t-maps were computed using identical procedures employed for fUSI. However, to account for the larger fMRI sample size, a more stringent statistical threshold (t = 11) was applied. The resulting fMRI CAPs displayed spatial patterns consistent with previously reported rsfMRI CAPs ^27^. Cross-modal correspondence of CAP topographies was quantified by computing voxel-wise Pearson correlations between unthresholded CAP t-maps derived from fUSI and fMRI. Inter-CAP spatial correlation matrices were computed independently for fUSI and fMRI CAPs (Fig. S12). In addition, a cross-modal fUSI–fMRI CAP correlation matrix was computed to highlight differences in CAP correspondence across modalities. CAP dynamics in the fMRI dataset were then characterized using the same transition-based framework applied to the fUSI data. One-step and two-step transition probability matrices were computed following removal of self-transitions, and transition graphs were thresholded using identical degree constrained thresholding (th_DC_) criteria.

## Acknowledgments

Funding: This work has been funded by the European Research Council (ERC) under the European Union’s Horizon 2020 research and innovation program ((#DISCONN; no. 802371 and no. 101125054 #BRAINAMICS to A.G.), and by the Brain and Machines Flagship Programme of the Italian Institute of Technology. A.G. is also supported by an endowment by Paolo and Sara Baracchino.

## Data and materials availability

All data needed to evaluate the conclusions in the paper are present in the paper and/or the Supplementary Materials. Raw fUSI data supporting the conclusions of this study are publicly available at https://zenodo.org/records/18486493. All analysis code is available on GitHub at https://github.com/functional-neuroimaging/PepeMariani_2026 and will be referenced by a DOI upon publication.

