## Supplementary Materials for "Structural and dynamic embedding of the mouse functional connectome revealed by functional ultrasound imaging (fUSI)"

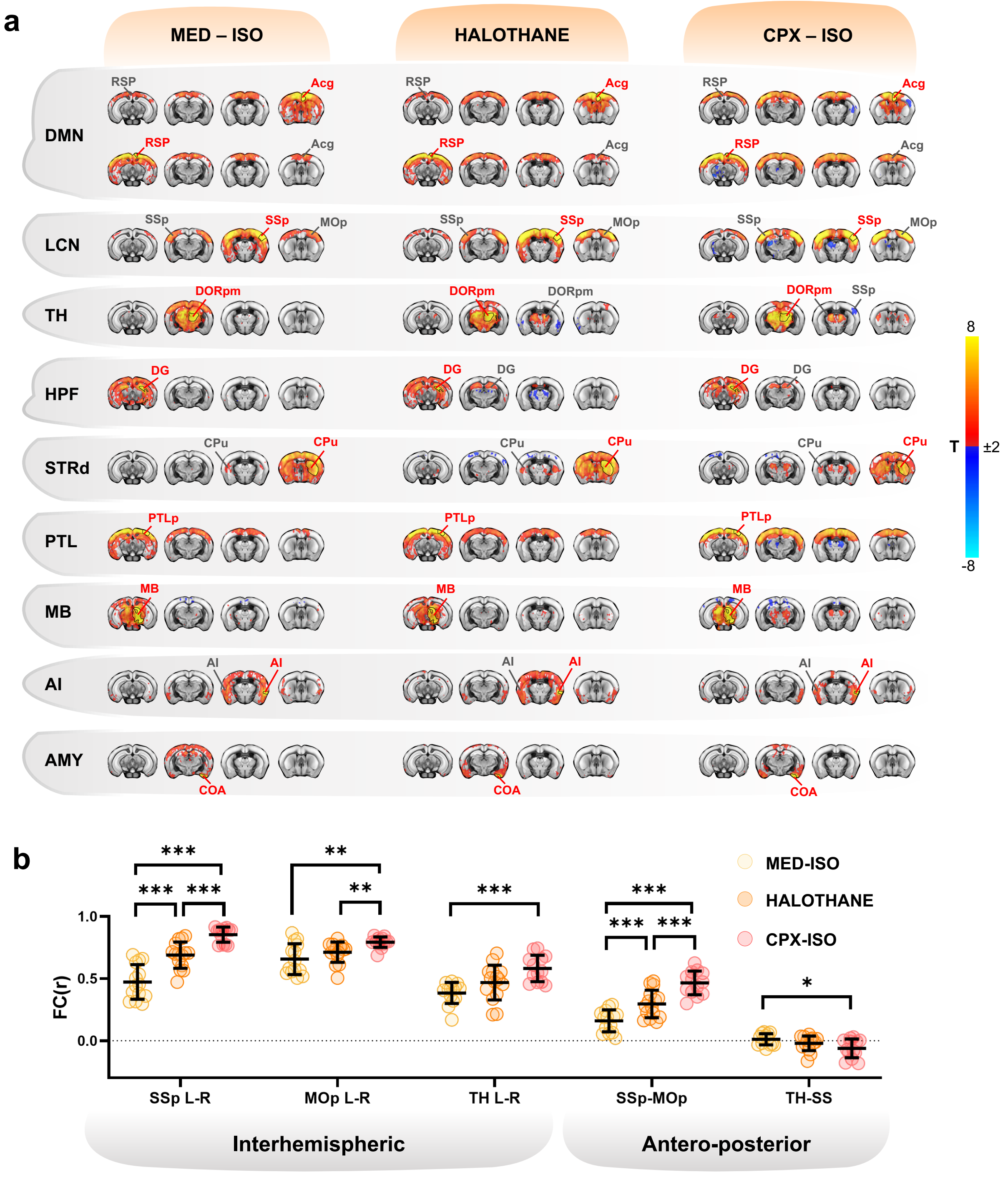

Figure S1. Resting-state fUSI network organization across light-sedation conditions. *(a) Seed-based connectivity maps for representative ROIs across three light-sedation groups (Med-Iso, halothane, Cpx-Iso). Networks shown include DMN, LCN, thalamic (TH), hippocampal formation (HPF), dorsal striatal (STRd), posterior parietal (PTL), midbrain (MB), insular (AI), and amygdalar (AMY). Group-level t maps are displayed (t > |2.1|, cluster-corrected, α = 0.05). Warm colors indicate positive t values, cool colors*

*negative. Seed location is indicated by red text labels.* ***(b)*** *Quantification of interhemispheric (SS L-R, MOp L-R, TH L-R) and anteroposterior (SS-MOp, TH-SS) functional connectivity across experimental groups. Statistics were performed on Fisher-transformed correlation coefficients (z values) using two-way repeated-measures ANOVA followed by Tukey's multiple-comparisons correction. Dots represent individual animals; data are shown as mean±SD. *p < 0.05, **p < 0.01, ***p < 0.001. ROI locations are indicated on the maps; abbreviations follow Allen Brain Institute nomenclature (Supplementary Table 1).* ***MOp,*** *primary motor cortex;* ***SSp,*** *primary somatosensory cortex;* ***TH,*** *polymodal thalamus; L, left; R, right.*

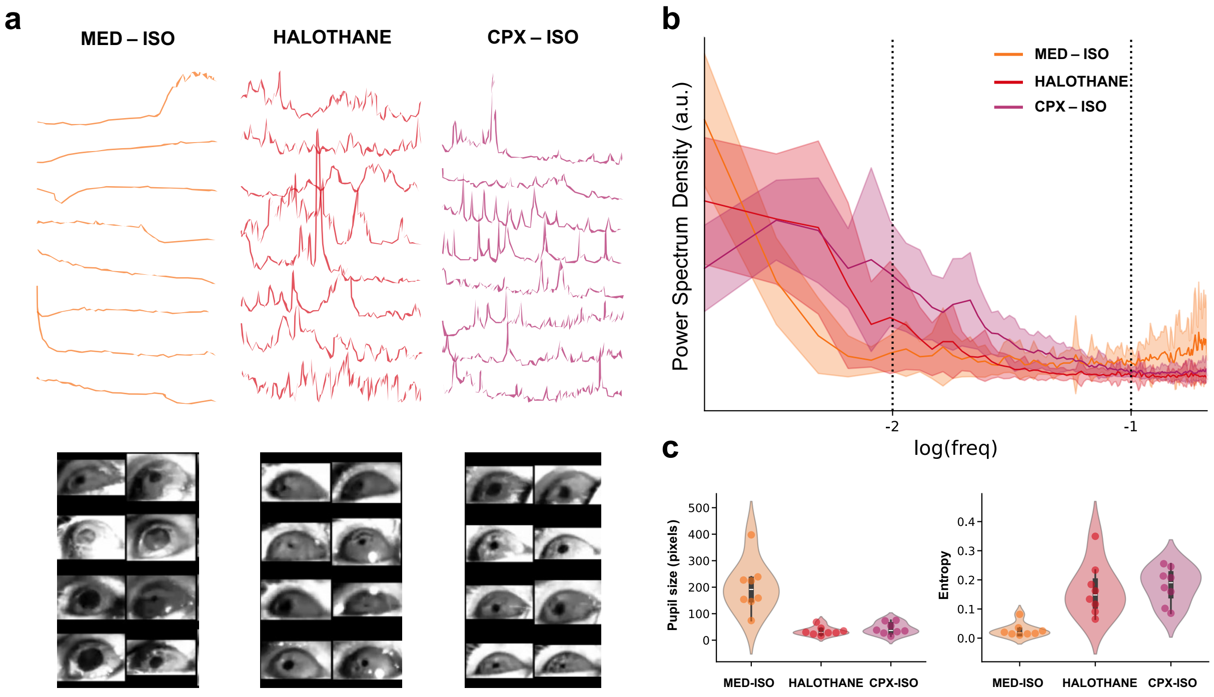

Figure S2. Pupil dynamics across light-sedation conditions. ***(a)*** *Representative pupil size time series (n = 8, each per group) under Med-Iso, halothane, and Cpx-Iso sedation (top), with corresponding pupil snapshots from the same recordings (bottom).* ***(b)*** *Semi-log power spectral density of pupil size fluctuations for each condition, normalized to unit area; dashed lines mark the resting-state functional connectivity band (0.01–0.1 Hz).* ***(c)*** *Group-level quantification of pupil dynamics. Left, median pupil size; right, temporal entropy of pupil time series. Dots represent individual animals; violins show group-wise distributions, with boxes indicating the interquartile range and the median (white line).*

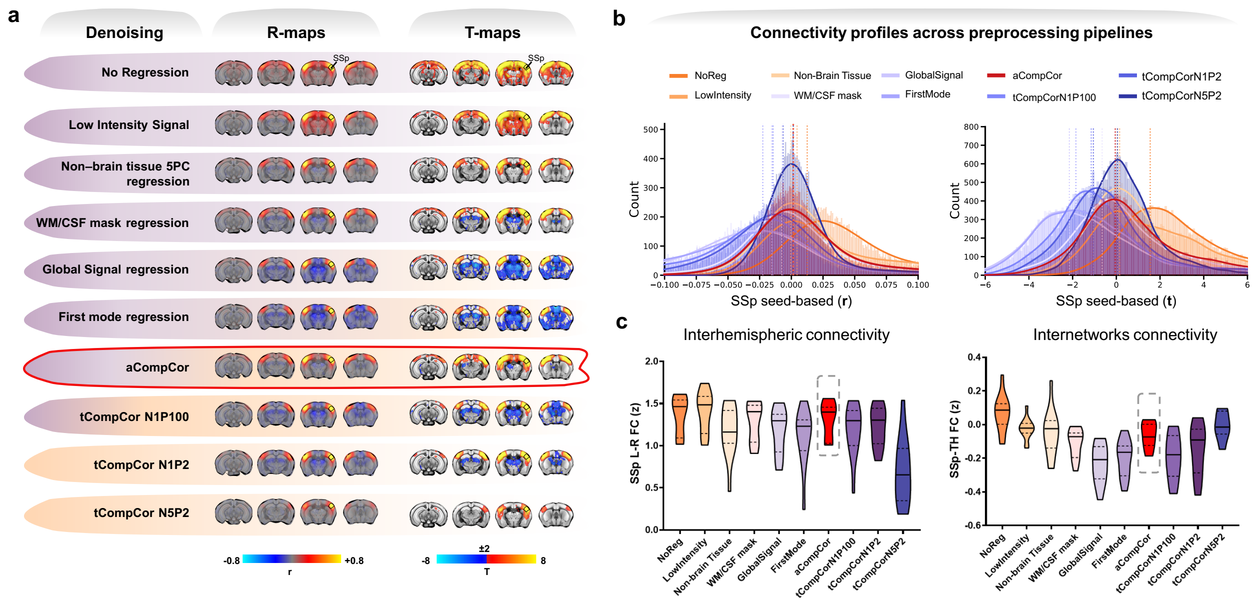

**Figure S3. Robustness of fUSI functional connectivity across preprocessing pipelines.** ***(a)*** *Seed-based connectivity maps for the primary somatosensory cortex (SSp) obtained using ten denoising/nuisance regression strategies. Unthresholded group-average correlation maps (r) and corresponding group-level t maps are shown; t maps are thresholded at t >|2.1| (cluster-corrected, α=0.05). The selected pipeline (aCompCor) is highlighted in red.* ***(b)*** *Distributions of connectivity values across preprocessing strategies, shown as correlation coefficients (r; left) and corresponding t values (right); vertical dashed lines indicate distribution maxima.* ***(c)*** *Pairwise connectivity across regression strategies. Left, interhemispheric connectivity between left and right SSp; right, internetwork connectivity between polymodal thalamus (TH) and SSp. Violins show group-wise distributions, with the median (solid line) and first and third quartiles (dotted lines). NoReg, no regression; LowIntensity, low-intensity signal regression; Non-Brain Tissue 5PC, regression of top 5 principal components from non-brain tissue; WM/CSF, white matter and cerebrospinal fluid regression; GlobalSignal, global signal regression; FirstMode, first-mode regression; aCompCor, anatomical CompCor; tCompCor N1P100/N1P2/N5P2, temporal CompCor variants (see Methods).*

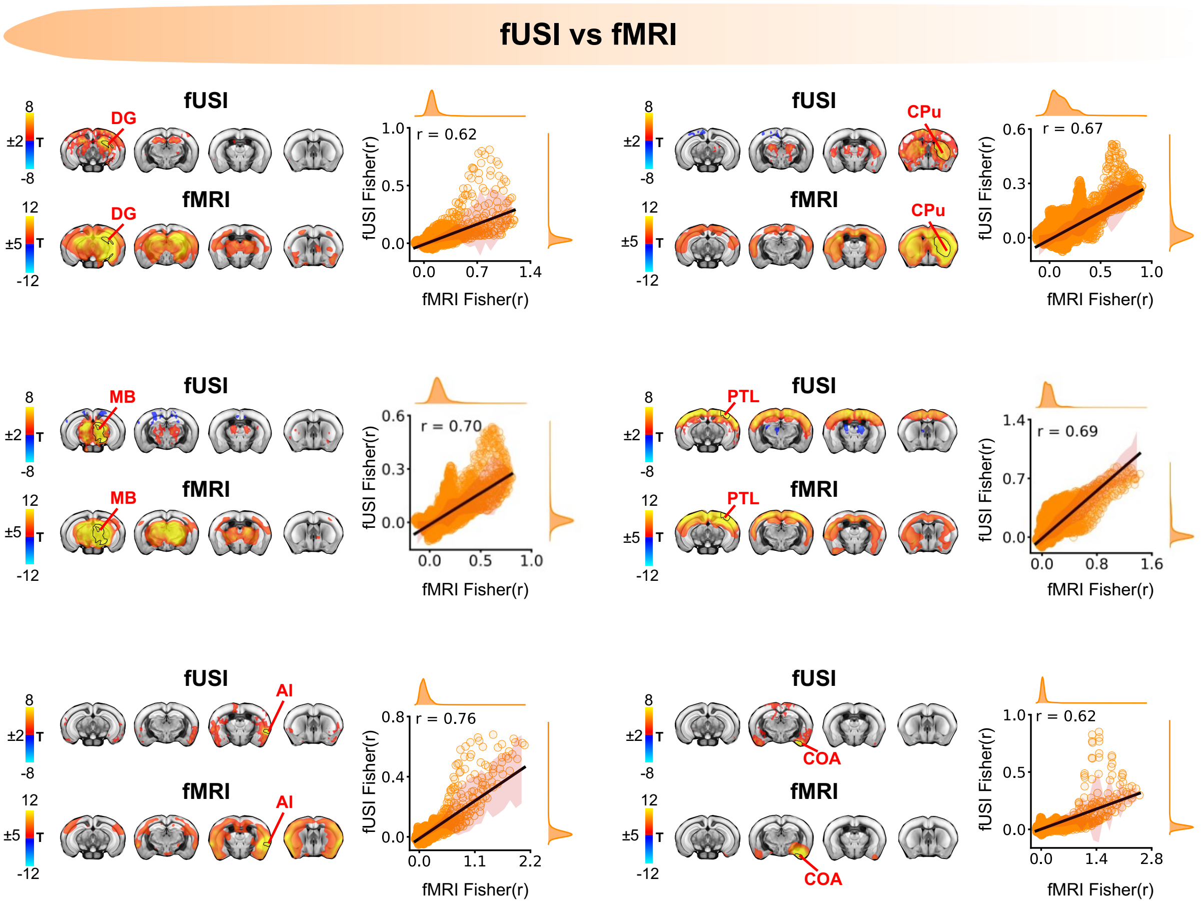

**Figure S4. Cross-modal correspondence of cortical and subcortical RSNs mapped with fUSI and fMRI.** *Seed-based connectivity maps for fUSI and fMRI for dentate gyrus (DG), caudate-putamen (CPu), midbrain (MB), posterior parietal cortex (PTL), insular cortex (AI), and cortical amygdalar area (COA) seeds. Seed location is indicated by red text labels. Corresponding voxel-wise spatial correlations between unthresholded Fisher-transformed connectivity maps from the two modalities are also shown. Solid lines indicate linear fits, shaded areas denote SD. Correlation coefficients (r) are reported for each seed. fUSI maps are shown as group-level t maps thresholded at t > |2.1| and fMRI maps at t > |5| (cluster-corrected, α = 0.05).*

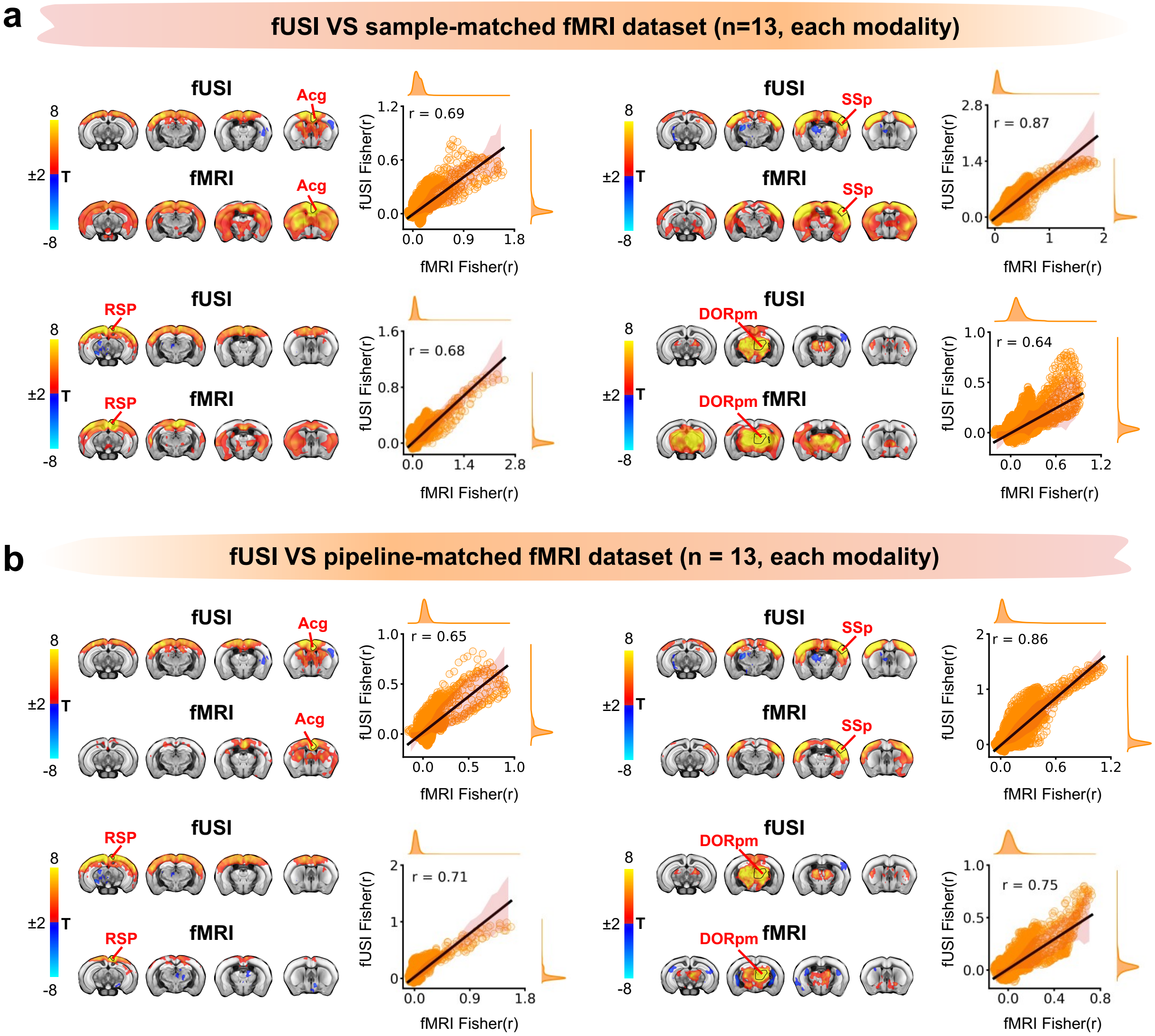

Figure S5. Robust cross-modal correspondence between fUSI and fMRI RSNs under matched sample size and preprocessing. ***(a)*** *fUSI-fMRI cross-modal comparison using a randomly selected sample-matched fMRI dataset (n = 13 animals per modality).* ***(b)*** *fUSI–fMRI comparison using a preprocessing-matched fMRI dataset (n = 13 per modality), processed with the same aCompCor-based denosing. For each seed (Acg, RSP, SSp, DORpm), group-level seed maps are shown as t maps thresholded at t > |2.1| (cluster-corrected, α = 0.05), and voxelwise correspondence is quantified as the spatial correlation between unthresholded Fisher-z connectivity maps (marginals are also shown). Seed locations are indicated on the maps in red. Acg, anterior cingulate cortex; RSP, retrosplenial cortex; SSp, primary somatosensory cortex; DORpm, polymodal thalamus.*

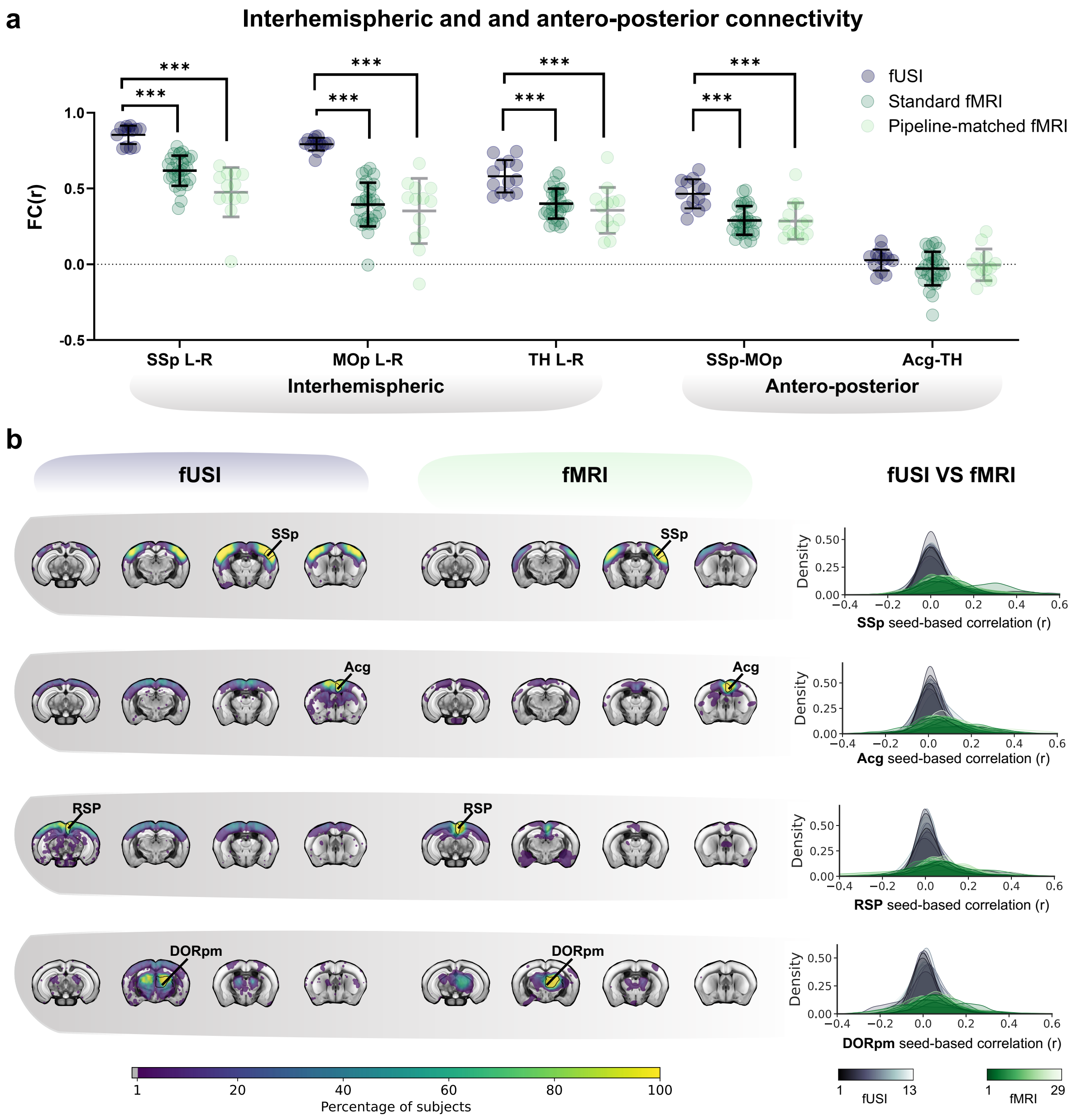

Figure S6. fUSI provides more stable FC mapping than fMRI. ***(a)*** *Interhemispheric and anteroposterior connectivity strength (r) across modalities. Functional connectivity was compared between fUSI (n=13) and a standard fMRI dataset (n = 29, standard preprocessing), and between fUSI and a pipeline-matched fMRI dataset (matched sample size, n = 13 for each modality; same aCompCor-based denosing). Metrics include interhemispheric coupling and antero-posterior coupling. Statistics were performed on Fisher-transformed correlations (z) using two-sample tests with Benjamini–Hochberg FDR correction (q = 0.05). Dots denote individual animals; ***p < 0.001.* ***(b)*** *Prevalence maps showing the percentage of subjects exhibiting suprathreshold (|z| > 2.5) seed-based connectivity for SSp, Acg, RSP and DORpm. Voxelwise density plots for fUSI (n = 13) and a standard fMRI dataset (n = 29). Cross-modal distributions were significantly different in all seeds (two-sided Kolmogorov–Smirnov test, p < 0.001). Acg, anterior cingulate cortex; DORpm, polymodal thalamus; MOp, primary motor cortex; RSP, retrosplenial cortex; SSp, primary somatosensory cortex; TH, polymodal thalamus. L, left; R, right.*

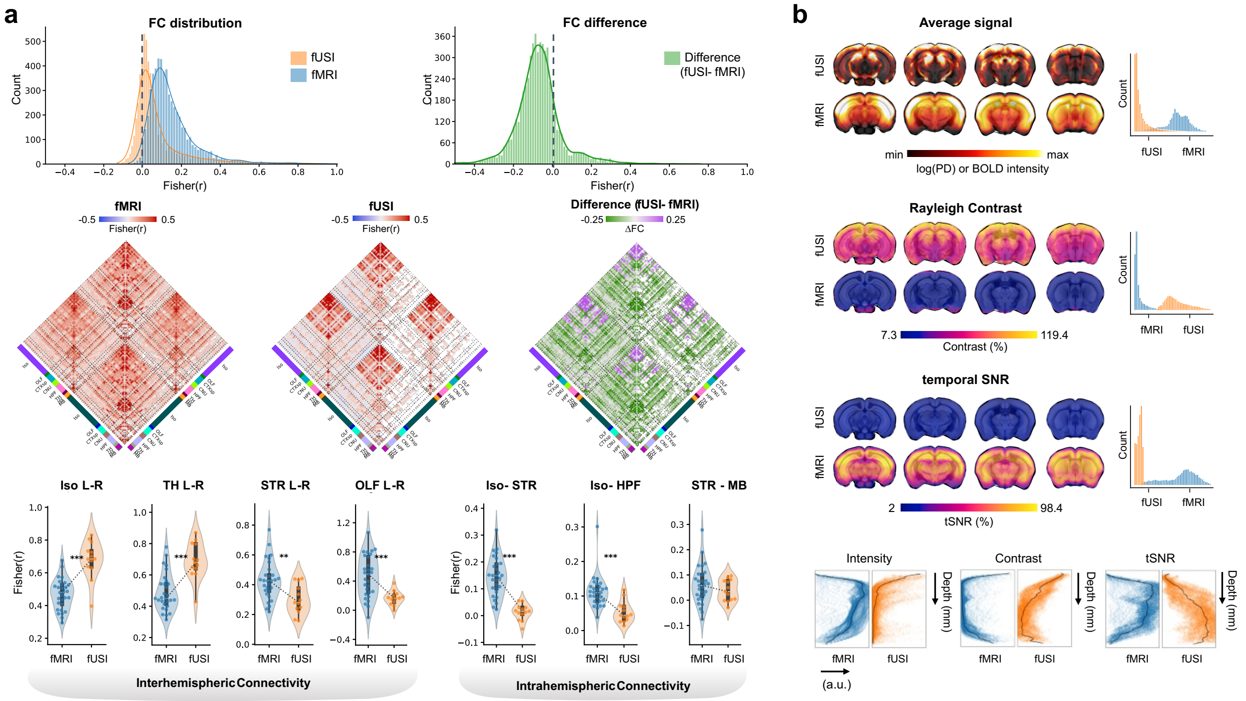

**Figure S7. Modality-dependent differences in functional connectivity and signal properties between fUSI and fMRI.** ***(a)*** *Comparison of functional connectivity (FC) estimates across modalities. Top: distributions of Fisher r-transformed FC values for fUSI and fMRI (left) and their difference (fUSI-fMRI; right). Middle: group-average FC matrices for fMRI, fUSI, and their difference, ordered by Allen Ontology macroregions and hemisphere. Bottom: Network-level connectivity quantifications are shown for representative interhemispheric and intrahemispheric pairs of regions.* ***(b)*** *Top: comparison of voxelwise signal properties between fUSI and fMRI. Population-average maps (fUSI, n = 13; fMRI, n = 29) and corresponding distributions are shown for average signal, Rayleigh contrast, and temporal signal-to-noise ratio (tSNR). Bottom: depth profiles of each metric along the dorsoventral axis, computed by averaging across the anteroposterior and mediolateral dimension. Iso, Isocortex; TH, Thalamus; STR, Striatum; OLF, Olfactory areas; HPF, Hippocampal formation; MB, Midbrain.*

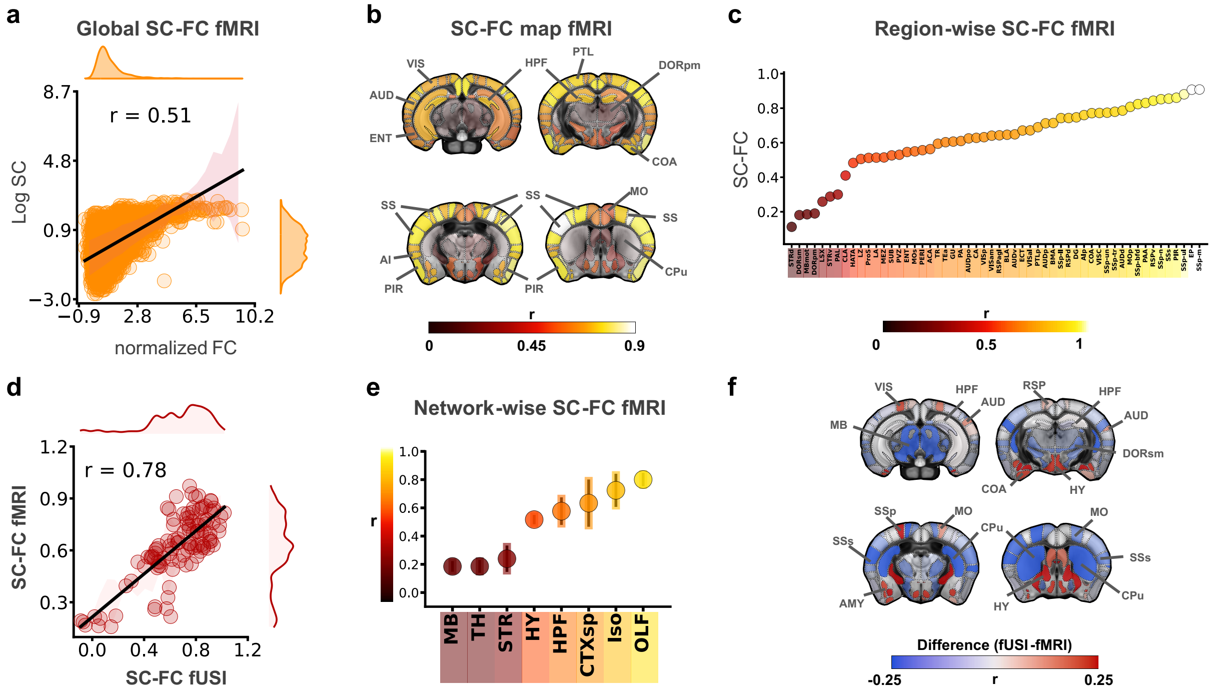

**Figure S8. Structure-function relationship in rsfMRI.** ***(a)*** *Global rsfMRI structure-function correspondence (SC-FC) across 108 regions of interest (ROIs).* ***(b)*** *Region-wise SC-FC correspondence in fMRI, computed as the correlation between structural and functional connectivity profiles for each ROI, projected onto the Allen Brain Atlas template. Color and transparency jointly encode correspondence strength.* ***(c)*** *Region-wise SC-FC values derived from (b). Dots represent correlation coefficients for 54 regions (averaged across hemispheres), ordered by increasing magnitude.* ***(d)*** *Cross-modal comparison of region-wise SC-FC profiles between fUSI and fMRI. Shaded area denotes SD. Marginal density plots are also reported.* ***(e)*** *Network-wise SC-FC in fMRI obtained by aggregating ROI-wise values within major Allen macroregions (means±SD across constituent ROIs).* ***(f)*** *Difference map of SC-FC correspondence between modalities (fUSI–fMRI). Full ROI nomenclature is reported in Supplementary table 1.*

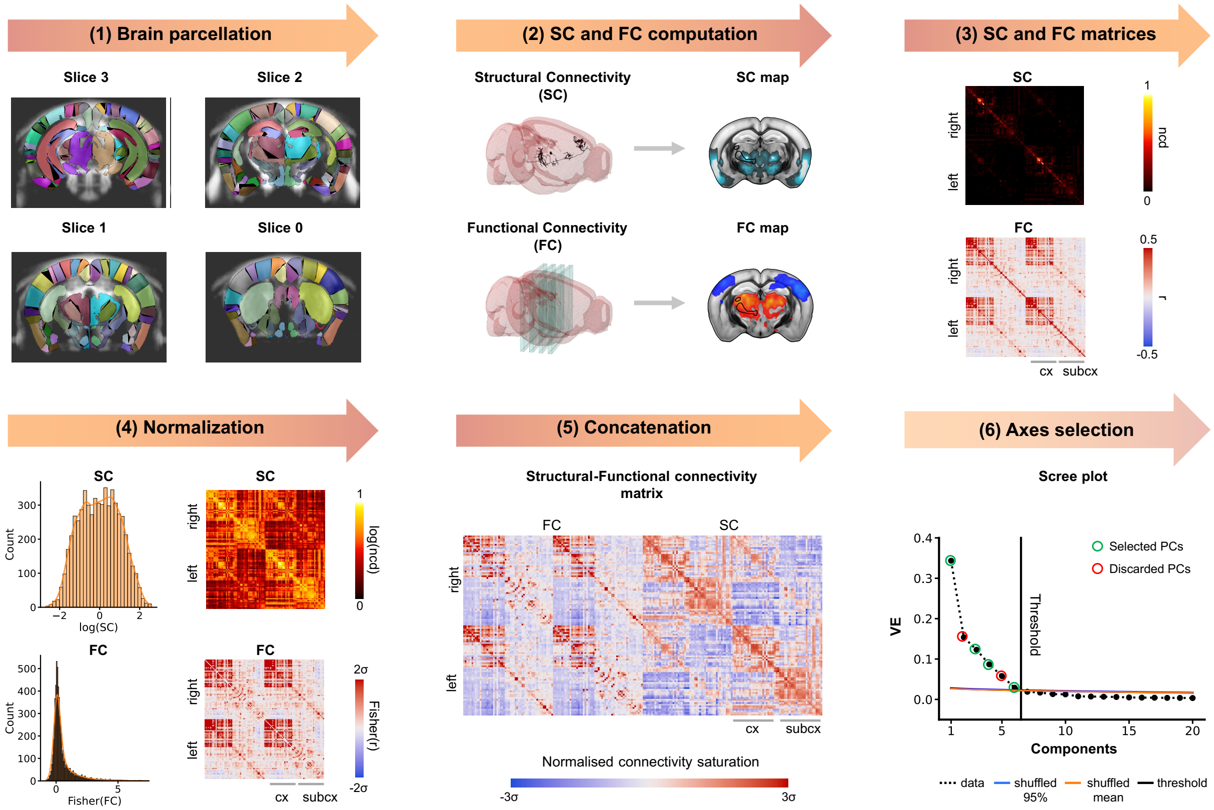

**Figure S9. Overview of the structural–functional connectivity (SC-FC) analysis workflow.** *(1) Brain parcellation into 108 unilateral regions of interest (ROIs) mapped onto the four fUSI imaging planes. (2,3) Structural connectivity (SC) was derived from the Allen Mouse Brain Connectivity Atlas and functional connectivity (FC) from resting-state fUSI, yielding ROI-by-ROI SC and FC matrices. Connectivity values were normalized (log-transformed SC, Fisher-transformed FC) (4) and concatenated to form a joint SC-FC matrix (5). Principal component analysis (PCA) was applied to the joint matrix; the scree plot shows explained variance per component (6), with significant components identified using bootstrap-based thresholds (black line). Mean (orange line) and 95$^{th}$ percentile (blue line) of shuffled distributions are shown for reference, selected axes are circled in green (see methods).*

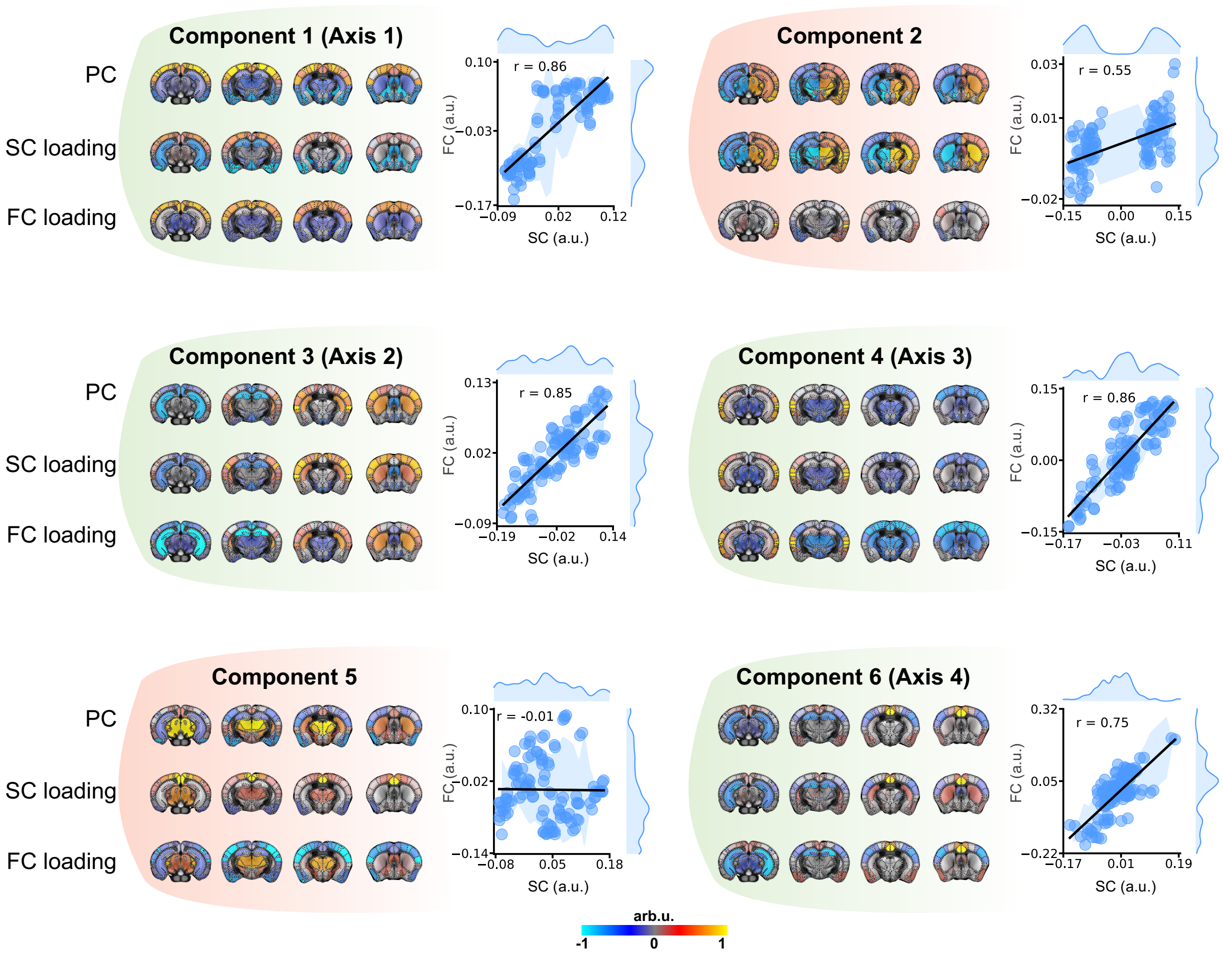

**Figure S10. Joint SC–FC principal components and modality loadings.** *Brain maps showing the spatial patterns of the first six joint SC-FC principal components (PCs), together with the corresponding ROI-wise structural (SC) and functional (FC) loadings. Dots represent individual regions of interest; solid lines indicate linear fits, and shaded areas denote SD. Marginal density plots show the distributions of SC (top) and FC (right) loadings. Components retained as joint axes (Components 1, 3, 4 and 6; green) show strong SC–FC alignment (r > 0.70), whereas discarded components (Components 2 and 5; red) show weaker correspondence.*

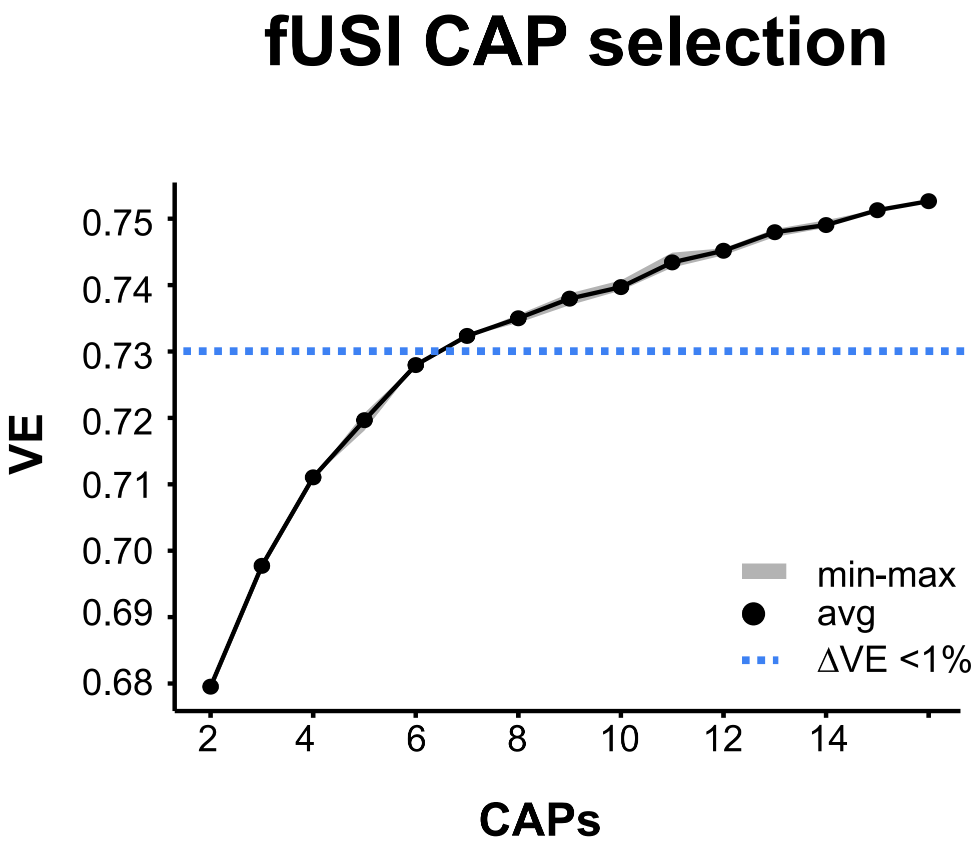

**Figure S11. CAP model-order selection.** *Variance explained (VE) as a function of the number of co-activation patterns (CAPs). Six CAPs were selected, accounting for ~73% of the variance; beyond this point, adding additional CAPs increased VE by <1% per extra component (ΔVE < 1%; blue dotted line). The shaded band indicates the min-max range across three independent runs.*

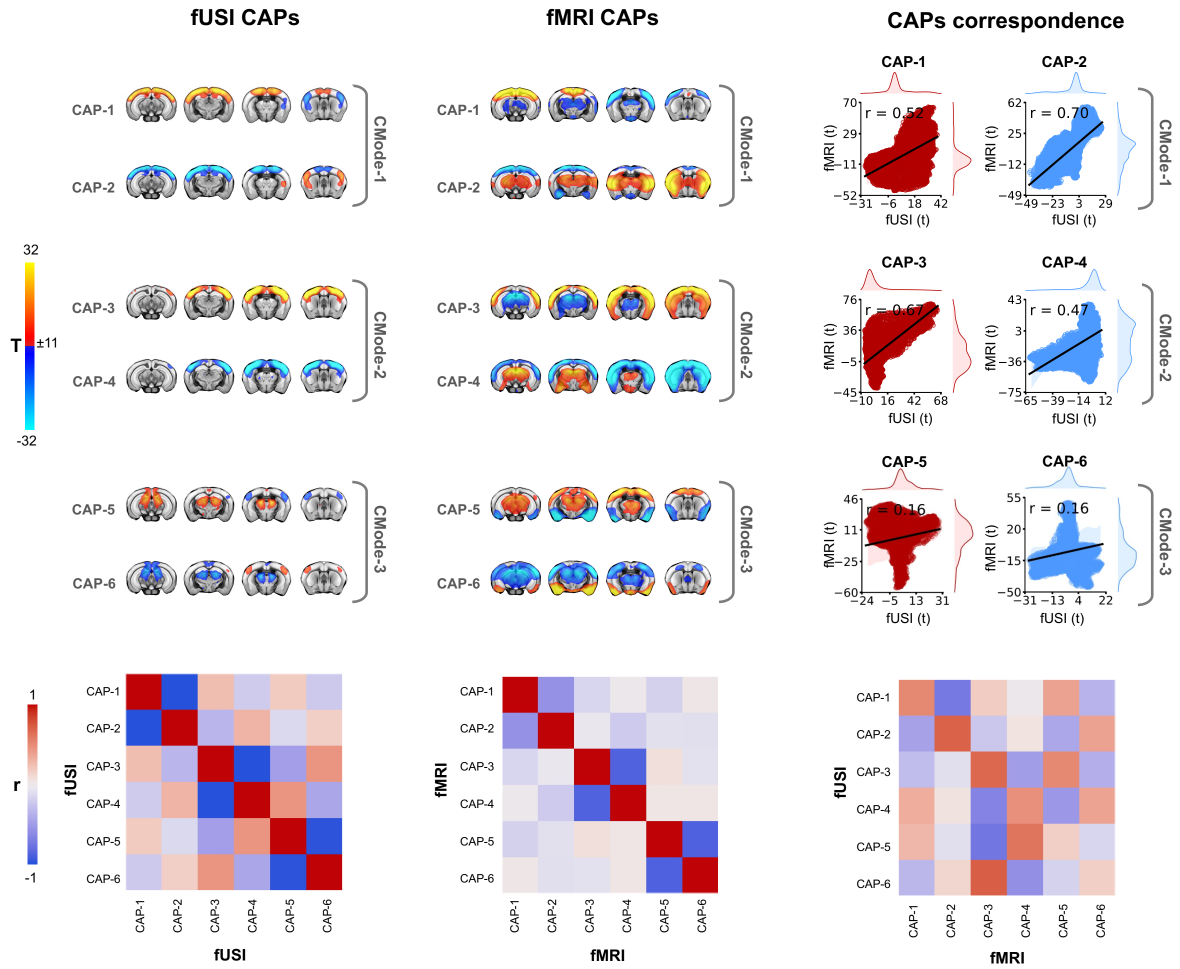

Figure S12. Cross-modal correspondence of co-activation patterns (CAPs) in fUSI and fMRI. *CAP maps for fUSI (left) and fMRI (middle) are grouped into three C-modes (C-mode 1: CAP1–2; C-mode 2: CAP3–4; C-mode 3: CAP5–6) and shown as group-level t maps (display thresholds: fUSI, t > 8; fMRI, t > 12; cluster-corrected). Right, cross-modal correspondence quantified as spatial correlations across voxels between unthresholded t maps from the two modalities for each CAP/antiCAP pair (red, CAPs; blue, antiCAPs); lines denote linear fits (shaded band, SD) with marginal density plots. Bottom, inter-CAP spatial correlation matrices computed within fUSI (left), within fMRI (center), and between fUSI and fMRI CAPs (right), illustrating modality-specific similarities and differences in CAP organization.*

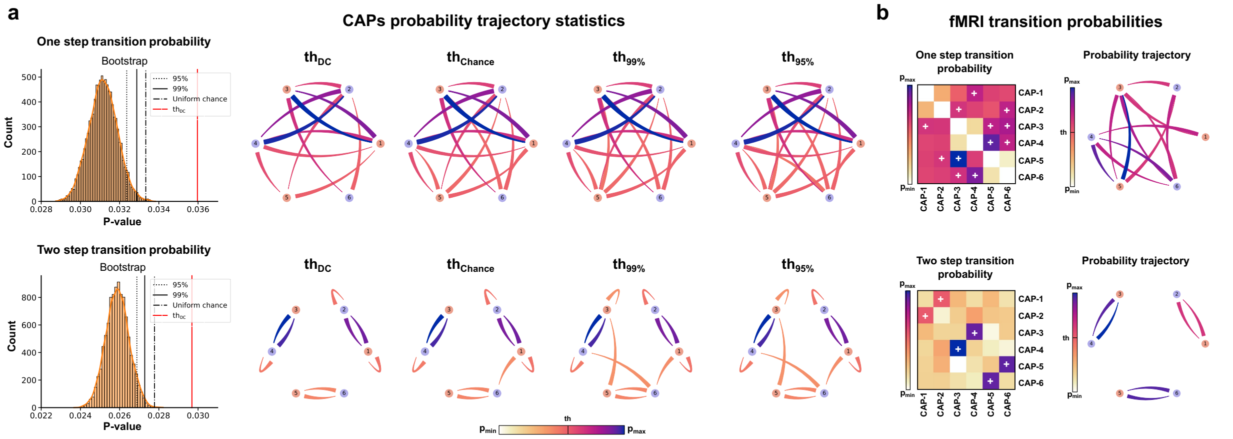

**Figure S13. Selection and validation of CAP transition thresholding.** ***(a)*** *Candidate thresholds for CAP transition graphs. Bootstrap/permutation-derived null distributions of one-step and two-step transition probabilities are shown (left), together with thresholds based on chance and percentile cutoffs (e.g., 95th and 99th) and degree-constrained (DC) thresholding. Directed transition graphs (right) illustrate how threshold stringency shapes the retained transition architecture.* ***(b)*** *One-step and two-step fMRI CAP transition probabilities thresholded using DC thresholding. Transition matrices and directed graphs show one-step and structured two-step trajectories linking opposing CAP–antiCAP pairs, consistent with attractor-like organization. (+) in matrices indicates suprathreshold edges.*

| Acronyms | Allen_ids | Parcel_ids | Networks | Alternative nomenclature |
|---|---|---|---|---|
| MOp | 985 | 1 | Iso | |
| MOs | 993 | 2 | Iso | |
| SSp-bfd | 329 | 3 | Iso | |
| SSp-ll | 337 | 4 | Iso | |
| SSp-ul | 369 | 5 | Iso | |
| SSp-tr | 361 | 6 | Iso | |
| SSp-un | 182305689 | 7 | Iso | |
| SSp-m | 345 | 8 | Iso | |
| SSp-n | 353 | 9 | Iso | SSp, SS |
| SSs | 378 | 10 | Iso | |
| VISC | 677 | 11 | Iso | |
| AUDd | 1011 | 12 | Iso | |
| AUDp | 1002 | 13 | Iso | |
| AUDpo | 1027 | 14 | Iso | |
| AUDv | 1018 | 15 | Iso | |
| VISal | 402 | 16 | Iso | |
| VISam | 394 | 17 | Iso | |
| VISp | 385 | 18 | Iso | |
| AIp | 111 | 19 | Iso | AI |
| RSPagl | 894 | 20 | Iso | |
| GU | 1057 | 21 | Iso | |
| ACA | 31 | 22 | Iso | Acg |
| RSPd | 879 | 23 | Iso | |
| RSPv | 886 | 24 | Iso | RSP |
| PTLp | 22 | 25 | Iso | PTL |
| TEa | 541 | 26 | Iso | |
| PERI | 922 | 27 | Iso | |
| ECT | 895 | 28 | Iso | |
| PIR | 961 | 29 | Iso | |
| COA | 631 | 30 | OLF | |
| PAA | 788 | 31 | OLF | |
| CA | 375 | 32 | HPF | |
| DG | 726 | 33 | HPF | |
| TR | 566 | 34 | HPF | |
| ENT | 909 | 35 | HPF | |
| SUB | 502 | 36 | HPF | |
| ProS | 484682470 | 37 | HPF | |
| HATA | 589508447 | 38 | HPF | |
| CLA | 583 | 39 | CTXsp | |

| | | | | |
|---|---|---|---|---|
| EP | 942 | 40 | CTXsp | |
| LA | 131 | 41 | CTXsp | |
| BLA | 295 | 42 | CTXsp | |
| BMA | 319 | 43 | CTXsp | |
| PA | 780 | 44 | CTXsp | |
| STRd | 485 | 45 | STR | CPu, CNU |
| STRv | 493 | 46 | STR | |
| LSX | 275 | 47 | STR | |
| PAL | 803 | 48 | STR | |
| DORsm | 864 | 49 | TH | |
| DORpm | 856 | 50 | TH | TH |
| PVZ | 157 | 51 | HY | |
| MEZ | 467 | 52 | HY | |
| LZ | 290 | 53 | HY | |
| MBmot | 323 | 54 | MB | |

Supplementary Table 1. ROI parcellation scheme. *List of all 108 ROIs used for parcellation following the Allen Mouse Brain Atlas ontology, including Allen structure IDs, parcel IDs used in this study, corresponding Allen macro-regions (network assignment), and alternative nomenclature used in the manuscript where applicable.*

| Subject_id | Session_name | Run | Included | Exclusion criterion |
|---|---|---|---|---|
| sub-cp230328a | ses-1CPTses1 | | Yes | |
| sub-cp230328b | ses-1CPTses1 | | Yes | |
| sub-cp230329a | ses-1MEDISOses1 | | Yes | |
| sub-cp230329b | ses-1CPTses2 | | Yes | |
| sub-cp230330a | ses-1CPTses3 | | Yes | |
| sub-cp230331a | ses-1CPTses4 | | Yes | |
| sub-cp230331b | ses-1CPTses4 | | Yes | |
| sub-cp230405a | ses-1MEDISOses2 | | Yes | |
| sub-cp230405b | ses-1MEDISOses2 | | Yes | |
| sub-cp230405c | ses-1MEDISOses2 | | Yes | |
| sub-cp230406a | ses-1HALOses1 | | No | Motion |
| sub-cp230406b | ses-1HALOses1 | run-1 | No | A better quality duplicate run was available |
| sub-cp230406b | ses-1HALOses1 | run-2 | Yes | |
| sub-cp230412a | ses-1CPTses5 | | Yes | |
| sub-cp230412b | ses-1CPTses5 | | No | Excessive Motion |
| sub-cp230412c | ses-1CPTses5 | | Yes | |
| sub-cp230413a | ses-1HALOses2 | | Yes | |
| sub-cp230413b | ses-1HALOses2 | | Yes | |
| sub-cp230413c | ses-1HALOses2 | | No | Excessive Motion |
| sub-cp230413d | ses-1CPTses6 | | Yes | |
| sub-cp230414a | ses-1MEDISOses3 | | Yes | |
| sub-cp230414b | ses-1MEDISOses3 | | Yes | |
| sub-cp230414c | ses-1MEDISOses3 | | Yes | |
| sub-cp230415a | ses-1CPTses7 | | Yes | |
| sub-cp230415b | ses-1CPTses7 | | Yes | |
| sub-cp230417a | ses-1HALOses3 | | Yes | |
| sub-cp230417b | ses-1CPTses8 | | Yes | |
| sub-cp230418a | ses-1HALOses4 | run-1 | No | A better quality duplicate run was available |
| sub-cp230418a | ses-1HALOses4 | run-2 | Yes | |
| sub-cp230419a | ses-1MEDISOses4 | | Yes | |
| sub-cp230420a | ses-1MEDISOses5 | | Yes | |
| sub-cp230421a | ses-1HALOses5 | | Yes | |
| sub-cp230424a | ses-1MEDISOses6 | | Yes | |
| sub-cp230424b | ses-1MEDISOses6 | | Yes | |
| sub-cp230504a | ses-1MEDISOses7 | | No | Excessive Motion |
| sub-cp230504b | ses-1MEDISOses7 | | Yes | |
| sub-cp230505a | ses-1HALOses6 | run-1 | No | Excessive Motion |
| sub-cp230505a | ses-1HALOses6 | run-2 | No | 3D registration failure |
| sub-cp230505b | ses-1HALOses6 | | Yes | |

| | | | |
|---|---|---|---|
| sub-cp230509a | ses-1HALOses7 | | Yes |
| sub-cp230509b | ses-1HALOses7 | | Yes |
| sub-cp230509c | ses-1HALOses7 | | Yes |
| sub-cp230510a | ses-1MEDISOses8 | | Yes |

**Supplementary Table 2. Dataset summary and exclusion criteria.** *All fUSI recordings included in the present dataset are listed. Columns indicate the subject ID, session identifier, run number (when applicable), whether the recording was included in the analyses, and the reason for exclusion.*
